# The acquisition of clinically relevant amoxicillin resistance in *Streptococcus pneumoniae* requires ordered horizontal gene transfer of four loci

**DOI:** 10.1101/2021.12.17.473165

**Authors:** Paddy S. Gibson, Evan Bexkens, Sylvia Zuber, Lauren A. Cowley, Jan-Willem Veening

## Abstract

Understanding how antimicrobial resistance spreads is critical for optimal application of new treatments. In the naturally competent human pathogen *Streptococcus pneumoniae*, resistance to β-lactam antibiotics is mediated by recombination events in genes encoding the target proteins, resulting in reduced drug binding affinity. However, for the front-line antibiotic amoxicillin, the exact mechanism of resistance still needs to be elucidated. Through successive rounds of transformation with genomic DNA from a clinically resistant isolate, we followed amoxicillin resistance development. Using whole genome sequencing, we showed that multiple recombination events occurred at different loci during one round of transformation. We found examples of non-contiguous recombination, and demonstrated that this could occur either through multiple D-loop formation from one donor DNA molecule, or by the integration of multiple DNA fragments. We also show that the final minimum inhibitory concentration (MIC) differs depending on recipient genome, explained by differences in the extent of recombination at key loci. Finally, through back transformations of mutant alleles and fluorescently labelled penicillin (bocillin-FL) binding assays, we confirm that *pbp1a*, *pbp2b*, *pbp2x*, and *murM* are the main resistance determinants for amoxicillin resistance, and that the order of allele uptake is important for successful resistance evolution. We conclude that recombination events are complex, and that this complexity contributes to the highly diverse genotypes of amoxicillin-resistant pneumococcal isolates.

## Introduction

Genomic plasticity through frequent and large-scale recombination drives the evolution of antibiotic resistance in *Streptococcus pneumoniae* (the pneumococcus) (Salvadori et al., 2019). This naturally competent member of the human nasopharyngeal commensal microbiota opportunistically causes otitis media as well as severe invasive diseases such as pneumonia, bacteremia, and meningitis (O’Brien et al., 2009; Shak et al., 2013). Despite conjugate vaccine introduction, the species remains an important human pathogen, as vaccine-escape and antibiotic resistant variants constantly arise (Croucher et al., 2011; Wyres et al., 2013).

Variants occur primarily through natural transformation and homologous recombination of DNA from strains or closely related species that occupy the same niche, such as *S. mitis* and *S. oralis* (Croucher et al., 2012; Dowson et al., 1993; Sibold et al., 1994). One case study showed more than 7% of the genome had been transferred between two pneumococcal strains during the polyclonal infection of a pediatric patient over three months (Hiller et al., 2010). Early studies of transformation in the pneumococcus showed uptake of fragments ranging from 2-6 Kb (Gurney and Fox, 1968), but cell-to-cell contact was found to facilitate transfer of long DNA fragments (Cowley et al., 2018), and recombinant regions as large as 30 – 50 Kb in length have been observed in clinically relevant lineages such as PMEN-1 (Croucher et al., 2011; Wyres et al., 2013, 2012). Events such as these have played a significant role in shaping the pneumococcal pangenome (Donati et al., 2010).

Natural competence is activated when the competence stimulating peptide (CSP) interacts with the ComDE two-component system (Straume et al., 2015) (Figure 1), where phosphorylated ComE activates transcription of the genes encoding the alternative sigma factor *comX* (Martin et al., 2013; Slager et al., 2019; Ween et al., 1999). ComX then activates transcription of the late *com* genes necessary for DNA uptake and recombination. This includes the type IV-like pilus (ComGC), endonuclease EndA, and transport proteins ComFA, and ComEC (Bergé et al., 2013; Lacks and Greenberg, 1976; Lacks and Neuberger, 1975; Laurenceau et al., 2013; Rosenthal and Lacks, 1980). RecA and the transformation-dedicated DNA processing protein A (DprA) are essential for single-stranded DNA (ssDNA) uptake without degradation (Attaiech et al., 2011). Although not fully understood, DprA and RecA are thought to polymerize on ssDNA upon cell entry, initiating the homology search, while additional DNA molecules are then bound by multiple SsbB proteins protecting them from degradation by endogenous nucleases (Attaiech et al., 2011; Morrison et al., 2007). It has been hypothesized that the accumulation of stable ssDNA-SsbB eclipse complexes in the cytoplasm enables successive recombination events during one competence window, increasing genetic plasticity (Attaiech et al., 2011). Sequestered ssDNA can then be accessed by the transformation-dedicated DNA processing protein A (DprA), which mediates the loading of RecA onto the pre-synaptic filament (Lisboa et al., 2014; Mortier-Barrière et al., 2007; Quevillon-Cheruel et al., 2012). DprA is also involved in competence shut-off, along with ComX and RpoD sigma factor competition, and CSP degradation by HtrA (Cassone et al., 2012; Martin et al., 2013; Mirouze et al., 2013; Piotrowski et al., 2009; Weng et al., 2013).

**Figure 1:**
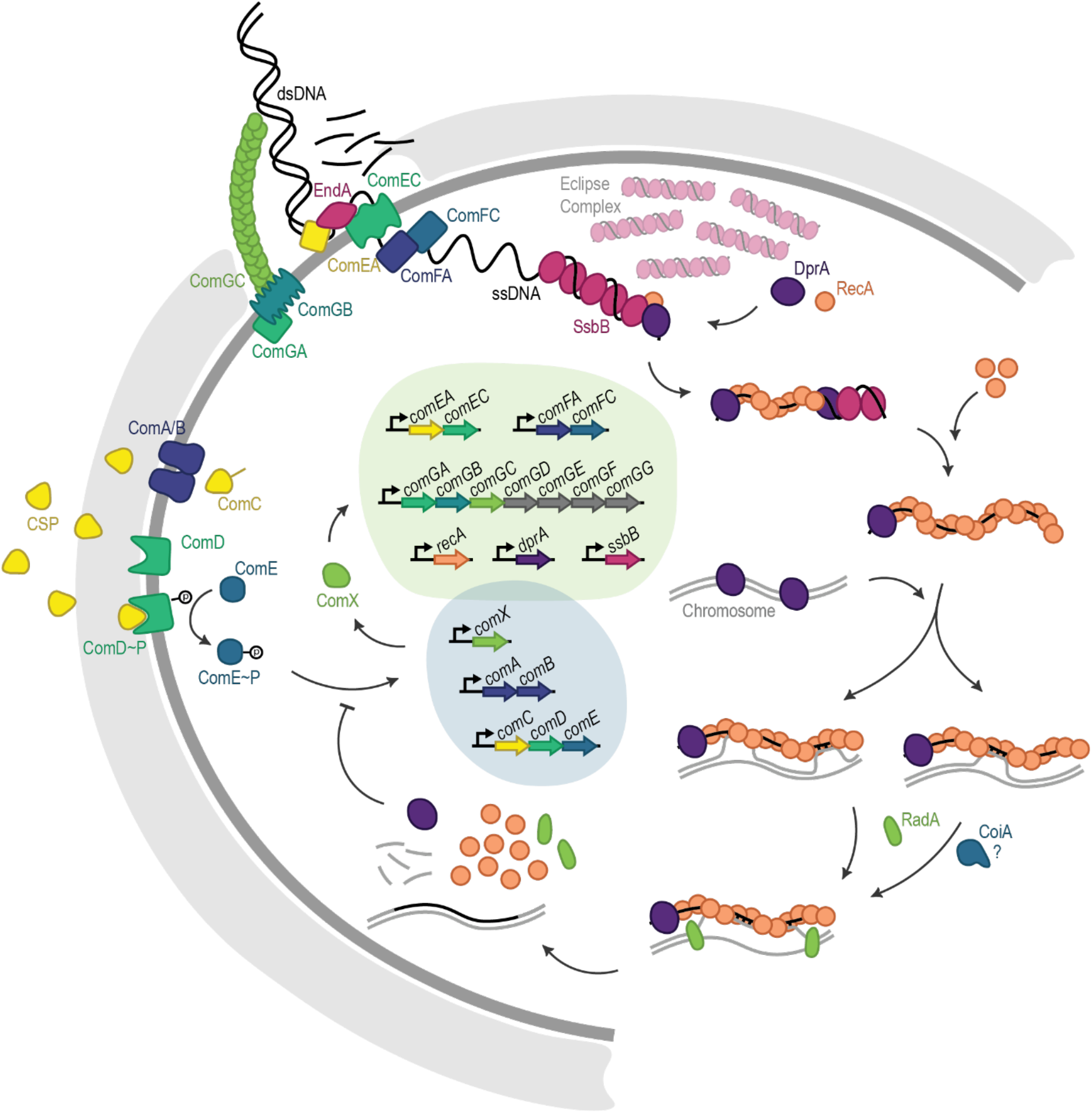
DNA uptake and recombination during natural competence in *Streptococcus pneumoniae*. Schematic overview of the regulatory steps required for the pneumococcus to uptake and recombine extracellular DNA. Competence is activated when CSP interacts with the ComDE two-component system, where phosphorylated ComE activates transcription of ComX and other early competence genes (darker grey bubble). This results in a positive feedback loop, where ComC is produced, then modified and exported from the cell by ComA/B as CSP. Meanwhile, ComX activates transcription of late competence genes, including the type IV-like pilus (ComGC), which brings DNA within reach of endonuclease EndA. EndA performs random nicks, then degrades one strand as the other is pulled into the cell and associated with DprA, RecA, and SsbB, to protect from degradation (exact mechanism not fully understood). DprA mediates loading of RecA onto SsbB-ssDNA eclipse complexes to form the presynaptic filament which then searches the chromosome for homology. RecA-ssDNA filaments may interact with the chromosome at multiple points, while successful base pairing between incoming and chromosomal DNA facilitates strand invasion and exchange. The helicase RadA facilitates D-loop extension, and CoiA is essential for successful recombination during transformation, although its exact role is not known. DprA is also involved in competence shut off, through blocking phosphorylation of ComE.

Competence is tightly controlled, with expression of most genes turned off within 20 minutes of CSP induction, providing a small window for DNA uptake and recombination (Alloing et al., 1998). To accelerate homology detection, the RecA-ssDNA presynaptic filament has been shown to interact with the chromosome in 3-dimensional space at multiple points of contact (Forget and Kowalczykowski, 2012; Wiktor et al., 2021; Yang et al., 2020). Displacement-loop (D-loop) formation is facilitated by base pairing between the chromosome and the ssDNA molecule, within the two DNA-binding sites of RecA (Bell and Kowalczykowski, 2016; Torres et al., 2019). Eight consecutive bases must be successfully paired before the DNA-DNA interaction is stabilized and strand exchange can occur (Hsieh et al., 1992; Mazin and Kowalczykowski, 1996). In *S. pneumoniae*, D-loop extension is promoted by RadA (DnaB-type helicase) which helps to unwind chromosomal DNA, extending recombination over long genomic distances (Marie et al., 2017), and while its function is not known, CoiA is essential for successful recombination (Desai and Morrison, 2007; Mortier-Barrière et al., 2007). Although initial testing is highly stringent, once crossover has started and the region of complementation is extended, stringency decreases, allowing some mismatches and thereby increasing desirable genetic variation (Carrasco et al., 2016; Danilowicz et al., 2015). This is aided by the generalized mismatch repair system in *S. pneumoniae* (hex) which is not induced during competence (Claverys and Lacks, 1986), is only able to identify mismatches in the donor-recipient heteroduplex before D-loop resolution (Claverys and Lacks, 1986; Gasc et al., 1989), (Gasc et al., 1989) and is easily overwhelmed by excess mismatches (Humbert et al., 1995).

Penicillin binding proteins (PBPs) are key β-lactam resistance determinants and are essential for bacterial cell wall synthesis by catalyzing the transglycosylation (TG) and transpeptidation (TP) reactions responsible for peptidoglycan (PGN) formation (Scheffers and Pinho, 2005). While highly conserved in penicillin (PEN) sensitive strains, PBPs of resistant isolates are highly variable, with up to 10% of amino acid changes (Hakenbeck et al., 2012). Significant amounts of this variation was recently confirmed to be acquired from *S. mitis* (Kalizang’oma et al., 2021). These large-scale changes alter the structure of the active sites, reducing β-lactam binding affinity (Hakenbeck et al., 2012). Amoxicillin (AMX) is the first line oral antibiotic prescribed for bacterial lower respiratory tract infections. AMX resistant strains first appeared in the mid-90s in the United States and France (Doern et al., 1996; Doit et al., 1999) and are currently on the rise in Spain, a phenomenon which correlates with increased usage of oral AMX/clavulanic acid (Càmara et al., 2018). Early AMX resistant lineages were closely related to known PEN resistance clones already circulating in the population (Doit et al., 1999; Stanhope et al., 2007). Sequencing of *pbp* alleles from susceptible and resistant strains found high variation in *pbp2x* and *pbp1a*, while SNPs shared between AMX resistant isolates were only found in *pbp2b* (Cafini et al., 2006; Chesnel et al., 2005; du Plessis et al., 2002; Kosowska et al., 2004). However, no SNP or block of SNPs in any one *pbp*, when transferred to a susceptible strain, has been sufficient to confer resistance in a susceptible strain (Hakenbeck et al., 1998; Kosowska et al., 2004). This is not entirely surprising, as the transpeptidase activity of PBPs is an essential function, and modifications to the active site are likely to have deleterious effects which need to be compensated for (Albarracín Orio et al., 2011)(Albarracín Orio et al., 2011). One mechanism of fitness compensation appears to be through substitutions in MurM, which results in abnormally high proportions of branched muropeptides in the cell wall (Filipe et al., 2002; Filipe and Tomasz, 2000; Garcia-Bustos and Tomasz, 1990; Smith and Klugman, 2001). Branched muropeptides were shown to have increased importance in strains depleted for Pbp2b, indicating a link between muropeptide composition and cell elongation (Berg et al., 2013). Other non-target proteins which have been implicated in resistance include CpoA (LafB) (Grebe et al., 1997) and CiaH (Guenzi et al., 1994).

Although multiple residues in Pbp1a, Pbp2x, and Pbp2b were found to be under positive selection in AMX resistant isolates (Stanhope et al., 2008), a specific a block of ten substitutions in the 590-641 region of the Pbp2b TP domain is repeatedly associated with AMX resistance (Cafini et al., 2006; du Plessis et al., 2002; Kosowska et al., 2004). However, transformation of *pbp2x*, *pbp2b*, and *pbp1a* alleles to AMX susceptible strain R6 did not recreate the AMX resistance of the donor strain, and the addition of *murM* had no effect. Only when genomic DNA was used could the original resistance level be achieved, indicating the role of a non-*pbp* or *murM* resistance determinant (Chesnel et al., 2005; du Plessis et al., 2002), or unknown epistatic interactions (Arnold et al., 2018). Interestingly, other studies have found a correlation between *murM* mutations and AMX resistance, although their presence depended on the *pbp2b* allele (Cafini et al., 2006; Càmara et al., 2018). Taken together, these studies strongly suggest the existence of multiple routes to AMX resistance and imply a tight link in the roles of Pbp2b and MurM in the mechanisms.

To study recombination and the development of AMX resistance in the pneumococcus in more detail and identify possible evolutionary routes towards AMX resistance, we sequentially transformed genomic DNA from a resistant clinical isolate (serotype 11A, ST6521, German National Reference Centre for Streptococci 2017, SN75752) into two different recipient strains coming from different clonal complexes (serotype 2 D39V and serotype 4 TIGR4). Whole genome sequencing showed recipient-dependent, genome-wide mutation uptake. In addition to identifying the necessary determinants for AMX resistance, we found a strong tendency towards the order of *pbp* and *murM* allele uptake. Long read Pacbio sequencing excluded a role for DNA methylation or genomic rearrangements or movement of genetic mobile elements as resistance determinants. We also investigated non-contiguous recombination, previously observed in *S. pneumoniae*, *Haemophilus influenzae*, and *Helicobacter pylori* (Croucher et al., 2012; Kulick et al., 2008; Mell et al., 2011), and propose a model for how these complex events occur. To conclude, we show that the uptake of four alleles from a resistant strain (*pbp2X*, *pbp2B*, *pbp1A* and *murM*), in a specific order, is required and sufficient for clinically relevant resistance development towards amoxicillin in sensitive *S. pneumoniae*.

## Results

### Serial transformation with genomic DNA from an AMX resistant strain increased AMX MIC

In order to understand the sequence of recombination events leading to the development of AMX resistance, a susceptible *S. pneumoniae* strain D39V (minimum inhibitory concentration (MIC) 0.01 µg/mL) was transformed in two successive rounds with genomic DNA originating from AMX resistant clinical isolate 11A (MIC 4 µg/mL) followed by selection on a range of AMX concentrations above MIC (Figure 2A, Table 1). In the first round, ten colonies were picked at random from the highest AMX concentration with growth (0.03 µg/mL AMX). The MICs of these strains ranged from 0.047 – 0.125 µg/mL, 4 – 10 times that of wild-type D39V (Figure 2B, Table S1).

**Figure 2:**
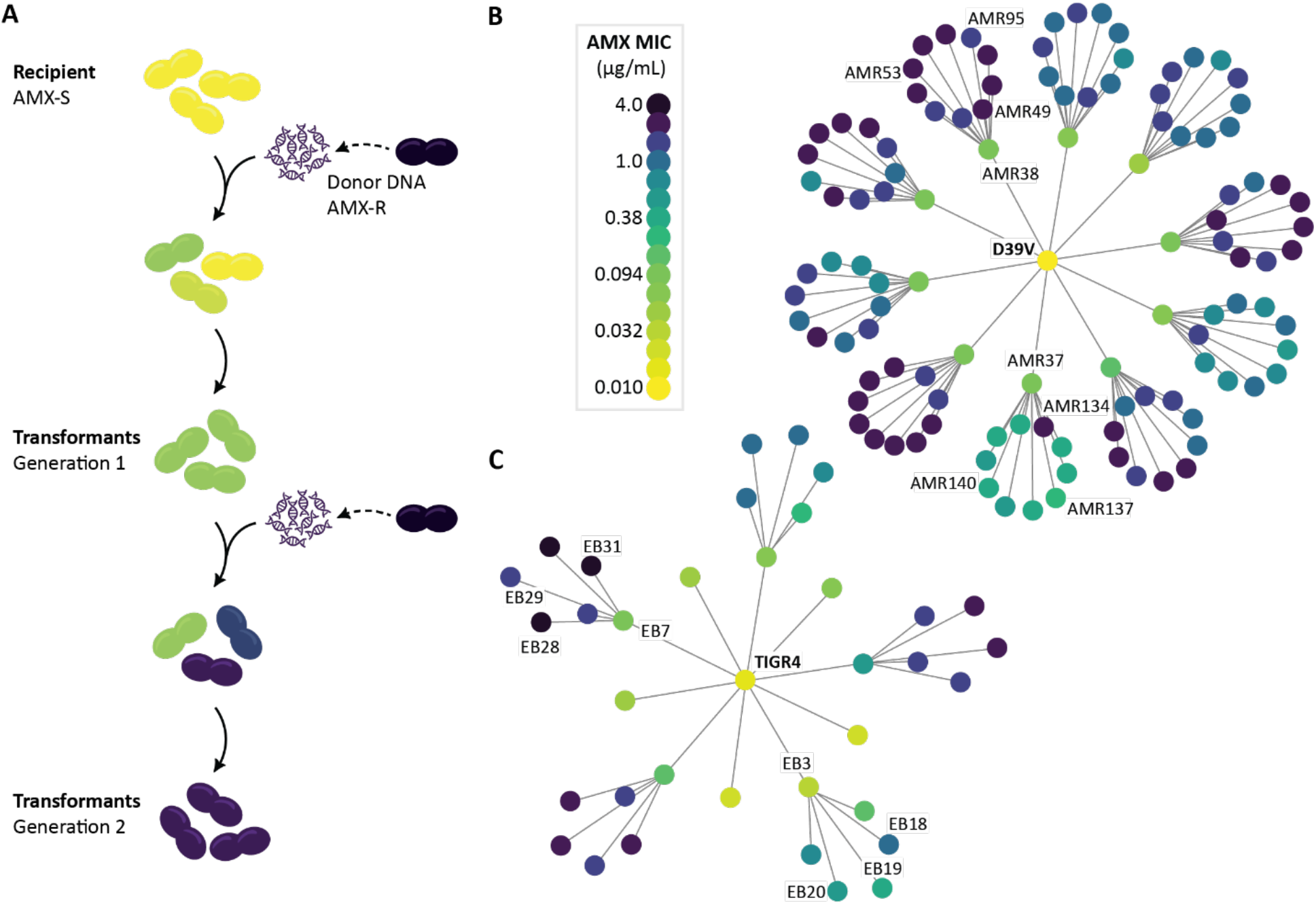
Transformation as an experimental tool to track the evolutionary path to AMX resistance. **(A)** Experimental design. Susceptible D39V and TIGR4 were transformed with genomic DNA from an AMX resistant clinical isolate then selected on a range of AMX concentrations. Ten colonies were picked at random from the highest concentration with growth (first round on 0.03 µg/mL and second round on 0.17 – 0.3 µg/mL, depending on MIC of the recipient strain). Isolated transformants with increased MICs were then subjected to a second round of transformation. **(B)** Pedigree chart of recombinant strains derived from D39V, and **(C)** from TIGR4. Color represents AMX MIC. Strains which were sequenced and used throughout the study are labelled.

**Table 1:** Key strains used in this study. The full collection of recombinant strains and their AMX MIC can be found in Table S1

| Strain Name | Strain ID | Genotype and relevant characteristics | AMX MIC (µg/mL) | reference |
| --- | --- | --- | --- | --- |
| D39V | VL1 | Serotype 2 laboratory strain, AMX susceptible | 0.001 | Slager et al., 2018 |
| TIGR4 | VL2177 | Serotype 4 laboratory strain, AMX susceptible | 0.016 | Tettelin et al., 2001 |
| 11A | SN75752<br>VL1313 | Serotype 11A ST6521 clinical isolate, AMX resistant | 4 | German National Reference Center for Streptococci |
| AMR37 | VL4481 | D39V; recombinant strain, generation 1 | 0.094 | This study |
| AMR38 | VL4482 | D39V; recombinant strain, generation 1 | 0.094 | This study |
| AMR53 | VL3196 | D39V; recombinant strain, generation 2 | 2 | This study |
| AMR95 | VL4484 | D39V; recombinant strain, generation 2 | 1.5 | This study |
| EB31 | VL4495 | TIGR4; recombinant strain, generation 2 | 4 | This study |
| D39V<sup>2x37</sup> | VL4496 | D39V; *pbp2x37* | 0.023 | This study |
| D39V<sup>2x38</sup> | VL4302 | D39V; *pbp2x38* | 0.064 | This study |
| D39V<sup>2b37</sup> | VL4497 | D39V; *pbp2b37* | 0.023 | This study |
| D39V<sup>2b38</sup> | VL4498 | D39V; *pbp2b38* | 0.023 | This study |
| D39V<sup>2x37-2b37</sup> | VL4499 | D39V; *pbp2x37*; *pbp2b37* | 0.094 | This study |
| D39V<sup>2x38-2b38</sup> | VL3895 | D39V; *pbp2x38*; *pbp2b38* | 0.094 | This study |
| D39V<sup>M53</sup> | VL3960 | D39V; *murM53* | 0.023 | This study |
| D39V<sup>1a53</sup> | VL4159 | D39V; *pbp1a53* | 0.016 | This study |
| D39V<sup>1a95</sup> | VL4160 | D39V; *pbp1a95* | 0.016 | This study |
| D39V<sup>2x38-2b38-M53</sup> | VL3961 | D39V; *pbp2x38*; *pbp2b38*; *murM53* | 0.38 | This study |
| D39V<sup>2x38-2b38-M95</sup> | VL3962 | D39V; *pbp2x38*; *pbp2b38*; *murM95* | 0.5 | This study |
| D39V<sup>2x38-2b38-1a53</sup> | VL3896 | D39V; *pbp2x38*; *pbp2b38*; *pbp1a53* | 0.75 | This study |
| D39V<sup>2x38-2b38-M53-1a53</sup> | VL3963 | D39V; *pbp2x38*; *pbp2b38*; *murM53*; *pbp1a53* | 2 | This study |
| D39V<sup>2x38-2b38-M95-1a95</sup> | VL4500 | D39V; *pbp2x38*; *pbp2b38*; *murM95*; *pbp1a95* | 1.5 | This study |
| AMR53<sup>2b31</sup> | VL4174 | AMR53; *pbp2b31* | 6 | This study |
| AMR95<sup>2b31</sup> | VL4184 | AMR95; *pbp2b31* | 6 | This study |

The second round of transformation resulted in 100 strains with MICs from 0.25 to 2 µg/mL (Figure 2B), lower than the donor strain, and a third round of transformation did not increase the MIC further. Interestingly, five of the ten lineages had seven or more strains with MICs of 1 µg/mL or higher. In contrast, the AMR37 lineage resulted in MICs from 0.25 – 0.5 µg/mL, with one exception at 2 µg/mL. These lineage-dependent differences in AMX MIC range (*p* = 5 x 10^-4^, Fisher’s Exact Test) suggested that mutations acquired in the first round contributed to the recombination events which occurred in the second round, and thus to the MICs of the resulting strains.

Similar lineage-dependent MIC patterns were seen when TIGR4 was used as the recipient strain (*p* = 1.00×10^-3^, Fisher’s Exact Test) (Figure 2B). Here, the AMX MICs of the first-generation ranged from 0.023 – 0.5 µg/mL, with one strain showing a more than 30-fold increase from wild-type TIGR4. In the second-generation, three recombinant strains reached the same MIC as the donor strain (4 µg/mL), suggesting potential key differences in the recipient strain genomes, or deleterious epistatic effects of AMX resistance mutations which affected resistance development through recombination.

### Cell lysis upon AMX treatment correlated with PBP affinities

To test whether transformed strains phenocopy the donor with regards to AMX resistance, phase contrast microscopy with an AMX concentration above the recipient MICs (1 μg/mL) was performed. A complete stall in growth was observed for both recipient strains and first-generation recombinant AMR38, with mild to severe lysis after 6 hrs of treatment (Figure 3A). In contrast, donor strain 11A grew to form a microcolony in this AMX concentration, while second-generation AMR53 grew more slowly and with abnormally elongated cells (Figure 3A). To test whether the increased resistance of AMR53 is related to alterations in the affinity of the PBPs towards AMX, we performed Bocillin-Fl labelling. Bocillin-Fl is a fluorescently labelled penicillin derivative that binds all six PBPs of *S. pneumoniae* (Zhao et al., 1999), and the intensity of labelling can be used as a proxy for the affinity of the PBP for β-lactam antibiotics (Kocaoglu et al., 2015). All six PBPs of D39V and TIGR4 had a high affinity for the fluorescently labelled penicillin (Figure 3B). Interestingly, TIGR4 showed some evidence of Pbp1A degradation products that bound Bocillin-Fl with high affinity even though it is 99.7 % identical to D39V Pbp1A. In contrast, 11A showed an altered PBP pattern, where PBP2b and Ppb1a showed complete loss of affinity, and Pbp2x was labelled more faintly and migrated more slowly (Figure 3B).

**Figure 3:**
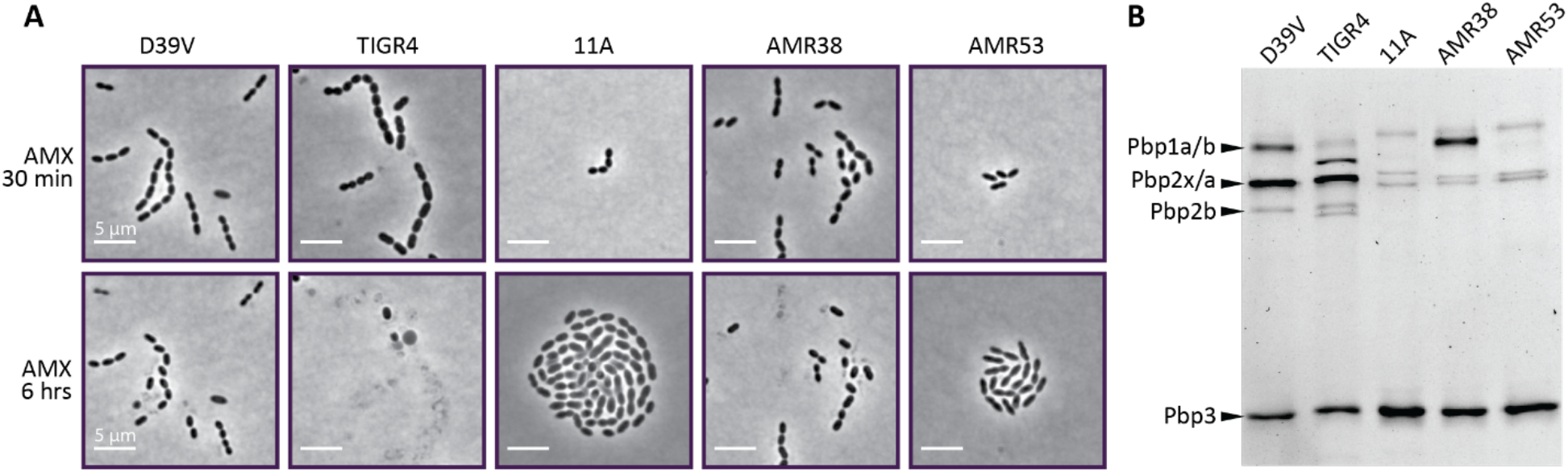
Phenotypic characterization of key strains used in this study. (**A**) Phase-contrast snapshots of recipient strains D39V and TIGR4, donor strain 11A, and recombinant strains AMR38 (generation 1) and AMR53 (generation 2) on C+Y agarose pads after 30 minutes and 6 hours of treatment with 1 µg/mL AMX. This concentration was above MIC for D39V, TIGR4, and AMR38, and lead to a complete stall in growth and partial lysis. Donor strain 11A grew well in this AMX concentration, although some heterogeneity in cell shape could be observed. Although lower than the AMX MIC for AMR53, this strain showed decreased growth and abnormally elongated cells, indicating cells were stressed. Scale bar is 5 µm. (B) Bocillin-FL binding patterns for the same five strains. Expected PBP band sizes for D39V are shown on the left. Image of Coomassie stained gel can be found in Figure S1)

Both first-and second-generation recombinants demonstrated a loss of affinity of Pbp2b and Pbp2x for Bocillin-FL, with PBP1a in AMR53 also showing reduced affinity, and potentially some degradation (Figure 3B). Together, these characterizations demonstrate that transformed recipients phenocopy the donor AMX resistant strain and have acquired PBPs with reduced affinity towards bocillin-FL.

### Recombination events clustered around resistance-associated loci

In order to examine the genetic differences in recombinant strains that may explain the differences in MIC, we selected eight recombinant strains from each experiment, and Illumina sequenced to ∼600-fold mean coverage. For the D39V experiment, we chose AMR38 and three of its descent strains, as this lineage had the highest MICs. To contrast, we also chose AMR37 and three descendent strains, which had the lowest and most variable MICs in the second generation (Fig 2B). We used a similar selection criteria for the TIGR4 experiment, taking strains from the high MIC EB7 lineage, and from the low MIC EB3 lineage (Fig. 2C). To be able to accurately map recipient to donor, we also performed long-read PacBio sequencing followed by short-read polishing on the 11A donor and TIGR4 recipient (an accurate genome map was already available for D39V (Slager et al., 2018)). Recombination was detected using SNPs, as described in Cowley et al. (2018). On average 0.38 % ± 0.22 % of the donor genome was transferred during a single transformation event, and the total percentage transferred after two rounds was 0.80 ± 0.05%. Although higher numbers of recombination events were detected in the D39V dataset, the average event length was larger in the TIGR4 dataset (D39V 0.79 Kb and TIGR4 2.29 Kb, Table S3 and S4), resulting in similar total amounts of donor genome transfer after two rounds of transformation (D39V 0.79 ± 0.23%, TIGR4 0.80 ± 0.16%, Table S3 and S4). Of note, the longest recorded recombination fragment was 4.00 Kb for D39V and 12.52 Kb for TIGR4.

Recombination events clustered around key loci for pneumococcal β-lactam resistance in both datasets; *pbp2x*, *pbp2b*, *pbp1a* and *murM* (Figure 4A and 5C). SNPs were also found at closely located loci, likely due to linkage. In the D39V strains, there was extensive recombination at distantly located sites. Examples include *ptvABC*, associated with vancomycin tolerance (Liu et al., 2017), and *murT*, which performs an amidation step essential for peptidoglycan crosslinking (Figueiredo et al., 2012; Münch et al., 2012).

**Figure 4:**
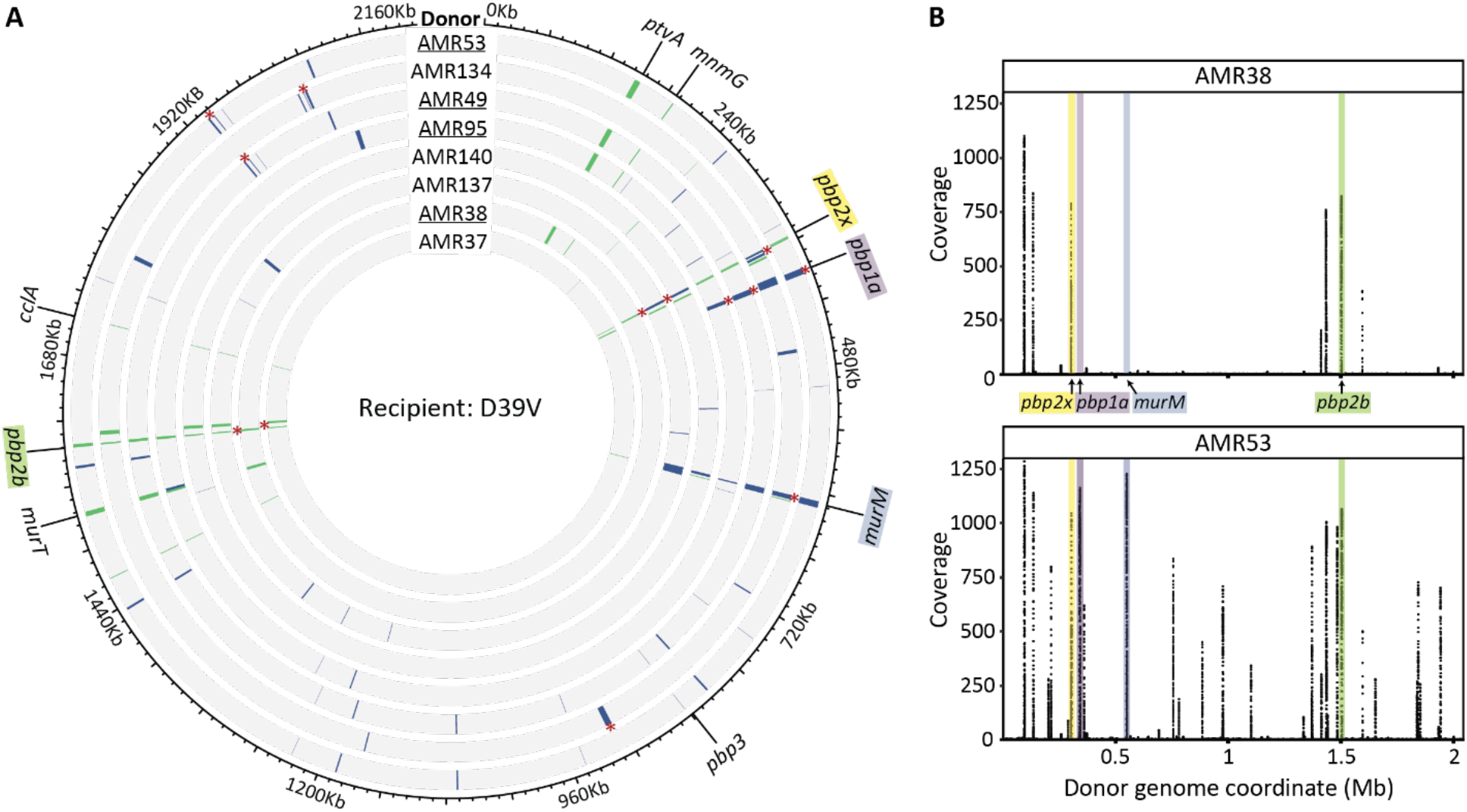
Widespread transformation and uptake of 11A donor DNA in recipient D39V upon AMX selection. **(A)** Circos plot with detected recombination events in D39V derived strains, arranged from lowest to highest MIC from the center. AMR38 and all derived strains are underlined, AMR37 and all derived strains are not. Grey indicates the sequences matches that of the recipient D39V, while recombination events are colored. Events acquired in the first round of transformation are shown in green, in the second round in blue, and lengths were increased to aid in visualization (see Table S3 for a list of all detected events). Recombination events determined to be non-contiguous are marked with (*). Non-contiguous events detected in the first-generation were removed from the data set on the second-generation strains, so are not marked. **(B)** Coverage of reads mapped competitively to the donor genome and plotted by coordinate beginning at the origin. Recombination events cluster at known β-lactam resistance determinants, *pbp2x*, *pbp2b*, *murM*, and *pbp1a*.

**Figure 5:**
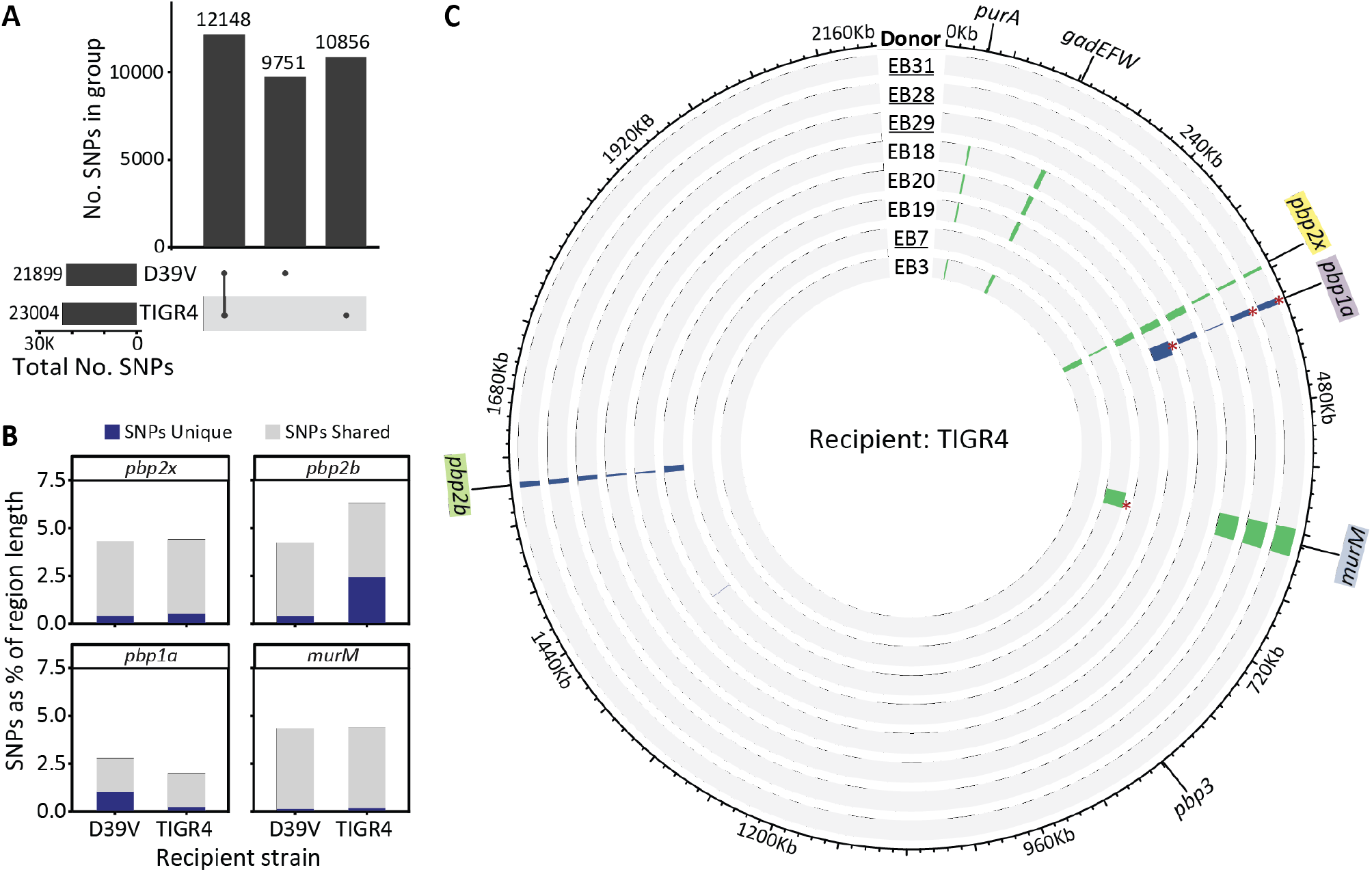
Uptake of four donor fragments is sufficient for AMX resistance in TIGR4. **(A)** Upset plot of shared and unique SNPs between each recipient and the donor. Bars on the left show the total number of SNPs for each recipient genome compared to the donor. In the central bar plot, from left to right, SNPs shared between D39V and TIGR4, SNPs found only in D39V, and SNPs found only in TIGR4. Although the two recipients differ from the donor strain by similar amounts of SNPs, only about half are unique. **(B)** SNP counts in pbp/murM genes, including 5 Kb up- and downstream. Number of SNPs unique to each recipient are shown in blue, and shared in light grey. **(C)** Circos plot showing detected recombination events in TIGR4 derived strains, arranged from lowest to highest MIC from inside to out. EB7 and all derived strains are underlined, EB3 and all derived strains are not. Grey indicates the sequences matches that of the recipient D39V, while recombination events are colored. Events acquired in the first round of transformation are shown in green, and in the second round in blue, and lengths were increased to aid in visualization (see Table S3 for a list of all detected events). Only six loci with recombination were identified in the entire data set, four of which are known β-lactam resistance determinants, *pbp2x*, *pbp2b*, *murM*, *pbp1a*.

In contrast, of the six total recombination loci in the TIGR4 dataset, only three were located distally from the *pbp* and *murM* genes, and all were detected based on the presence of very few SNPs.

We hypothesised that differences in recombination patterns could have arisen from reduced similarity with the donor genome, however comparing total shared and unique SNPs did not suggest major differences in variability between the two recipient genomes (Figure 5A). When considering variation around key loci, we saw that the TIGR4 *pbp2b* region was more divergent from the donor than D39V. The quality of donor DNA used may have also played a role. Together, these results demonstrate that regardless of the recipient strain, four loci are consistently acquired in high AMX resistant recombinants, namely *pbp2x*, *pbp2b*, *pbp1a* and *murM* (Figure 5B).

Structural variation and base modification detection with PacBio sequencing revealed no major structural changes or differences in methylated motifs in the D39V recombinant strains. However, a local rearrangement was identified at the *hsdS-creX* locus in EB7, EB28, and EB29. In this region, spontaneous reshuffling of *hsdS* genes by CreX results in different methyltransferase specificity of the type I restriction modification system SpnD39III (Feng et al., 2014; Manso et al., 2014; Slager et al., 2018). TIGR4 was found to be predominantly in the B-configuration, EB7 and EB29 in the D-configuration, and EB28 in the C-configuration, while all D39V-derived strains and 11A were mostly in the F-configuration. Differences in methylated motifs have transcriptomic consequences (Manso et al., 2014), however MIC differences among these recombinants can largely be explained by SNPs in key loci for β-lactam resistance.

### Non-contiguous recombination events likely result from a single donor molecule

Recombination is often thought of as a donor molecule forming a single D-loop at a homologous site on the recipient chromosome, which is then expanded in both directions until the requirement for homology is no longer met, at which point the interaction ends. However, we detected recombination events from one round of transformation located very close together on the chromosome, sometimes within the same gene but with separating recipient SNPs, a motif reminiscent of clinically resistant mosaic *pbp* alleles. This has also been noted previously in both *in vitro* pneumococcal and *H. influenzae* recombination studies and was hypothesized to result from a single donor molecule (Croucher et al., 2012; Mell et al., 2011; Mortier-Barriere et al., 1997). To determine if events were significantly close together, forming a single non-contiguous recombination event, a bootstrapping approach was applied to the distances between novel recombination events within the same strain (Croucher et al., 2012). In this approach, the shortest distances between all recombination events for all strains were pooled to form the test population. Bootstrap tests were performed 1000 times, where the test population was sampled with replacement, and the resulting distribution compared to the shortest recombination distance for each strain (*d_test_*). If *d_test_* was below the 0.05 quantile value in at least 95% of bootstrap tests, then the events were considered to be linked, and the next closest pairing was tested. We found 37 events across all strains that could be amalgamated into single events under the assumptions of this approach (Figure S2, Table S4).

In addition, using PCR amplified fragments of the *pbp2x-mraY* locus from 11A, we were able to test, for the first time, whether non-contiguous recombination events occurred more frequently when the template was provided as a single molecule, or as two overlapping fragments. By ligating additional regions of non-homologous DNA, we also tested the effect of molecule length on recombination.

Transformation efficiency increased with length of both the overall molecule, and the homologous region (Figure 6A). This is in line with previous studies showing the importance of molecule length and homology in pneumococcal transformation (Kurushima et al., 2020; Morrison and Guild, 1972). We hypothesize that increased length may increase the likelihood of spontaneous contact with the ComGC pilus, while also decreasing the likelihood of homologous region fragmentation by EndA, prior to entry (Morrison and Guild, 1973). The efficiencies of the short overlapping fragments were lower than that of the long fragments, although increased slightly when transformed in combination.

**Figure 6:**
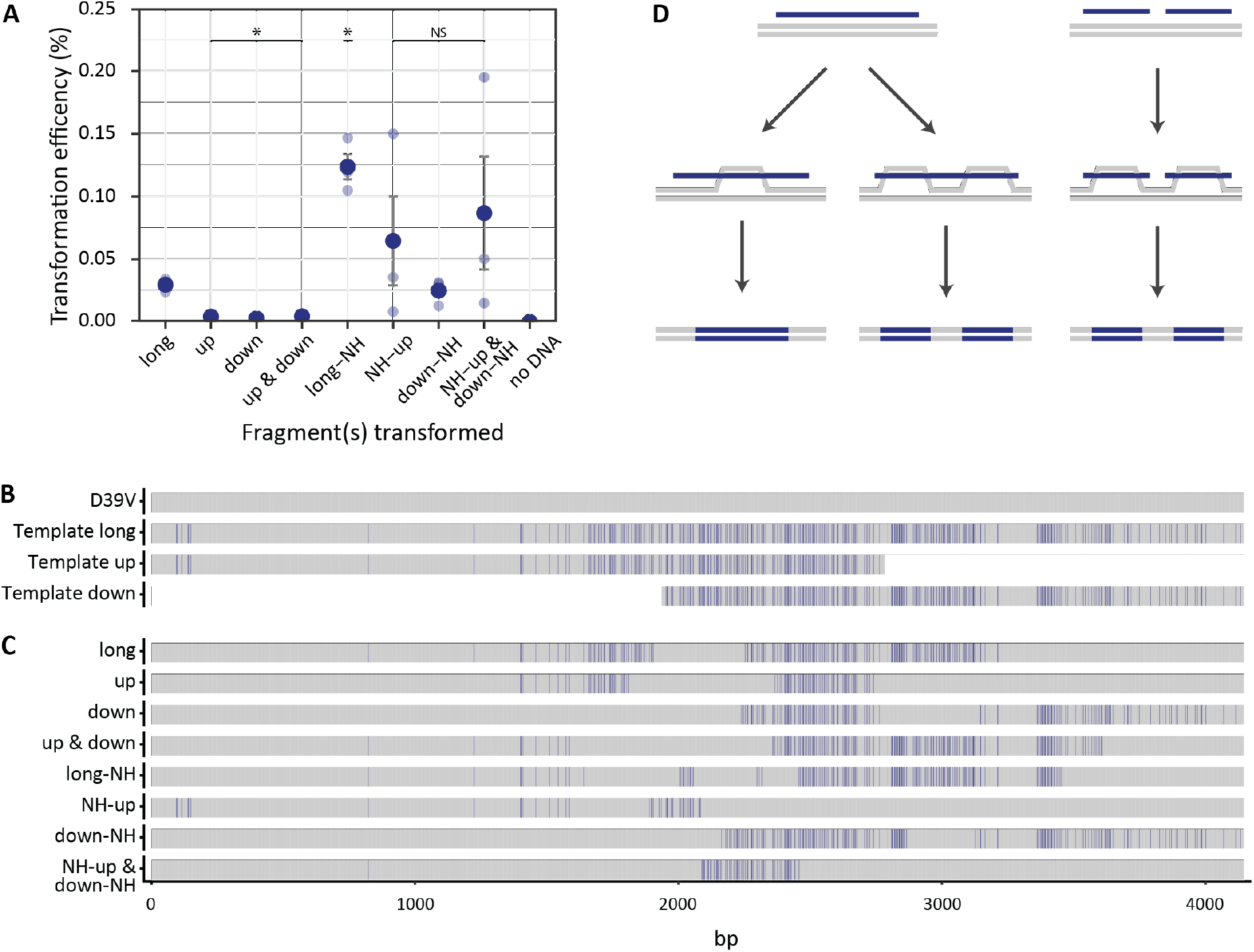
Fragmented transformation of donor DNA predominantly originates from non-contiguous recombination events. Three fragments were PCR amplified from the 11A strain *pbp2x* locus, one long 4.2 Kb fragment, and two overlapping short fragments (up 2837 bp, and down 2260 bp). In addition, the same fragments were ligated to stretches of non-homologous DNA, taking all total lengths to 6 Kb (long-NH, NH-up, and down-NH). **(A)** Efficiencies of transformation of fragments into D39V with selection on AMX (0.015 µg/mL). Statistically significant differences in efficiency compared to the long fragment was determined by *t* test with BH correction for multiple testing. NS not significant, * indicates *p* < 0.05. **(B)** Alignment of the homologous region of template fragments used in the experiment. Donor SNPs are shown in blue, while bases matching the recipient (D39V) are shown in grey. **(C)** Alignment of 4.2 Kb locus from colonies transformed with different fragments. Bases shared with donor but not with recipient are shown in blue. **(D)** Schematic showing hypothesized molecular mechanisms for non-contiguous and contiguous recombination. As shown in panel B, both events can take place, but based on transformation efficiencies, the process on the left with long donor DNA is more efficient.

Recombination occurred in all colonies sequenced. When two overlapping fragments were provided, uptake was often unequal, with either the upstream or downstream fragment incorporated more often. Recombination of both fragments was observed in 1/20 cases and was not observed when fragments were lengthened by non-homologous DNA (Figure S3A-H, Table S7).

Non-contiguous recombination was detected in all conditions, although the frequency of occurrence increased with length of homology (Figure 6C, Figure S3A-H, Table S7). It was observed both when the template was given as a single long fragment, and as two overlapping fragments.

Overall, the results suggested that the formation of multiple D-loops from either a single template molecule, or from multiple molecules, could lead to non-contiguous recombination events (Figure 6D). However, given that longer homology and total molecule length increased the efficiency of transformation partly by reducing strand degradation (Morrison and Guild, 1972), there may have been more opportunity for complex recombination patterns to occur when longer fragments were taken up.

### AMX resistant *pbp* and *murM* alleles are sufficient to explain AMX MICs

Substitutions in all four proteins (Pbp1a, Pbp2b, Pbp2x and MurM) were required to reach an MIC close or equal to that of the donor strain (Figure S4, and Table S5), but the order of uptake varied both stochastically, and depending on the recipient genome. The correlation between allele uptake and MIC increase was consistent in the D39V background. First-generation strains had mutations in *pbp2b* and *pbp2x*, the two essential PBPs for cell growth (Berg et al., 2013). In the second-generation, all strains acquired mutations in *murM*, and those with MICs higher than 1 µg/mL also had mutations in *pbp1a*. In TIGR4, *pbp2x* alleles were also acquired in the first-generation, however *pbp2b* mutations only appeared after the second round of transformation. Mutations in *murM* were recombined in EB7 in the first round of transformation, then were inherited vertically in that lineage, but were not acquired by EB3 or any descendant strains (Figure 5C). Although *murM* mutations were not present in all strains with *pbp1a* modifications, strains with recombination events at all four loci had higher MICs (Figure 2C). This indicated that *murM* mutations may be less important for successful integration and selection for *pbp1a* mutations but are key for expression of the AMX resistant phenotype of the donor (Filipe et al., 2002; Filipe and Tomasz, 2000; Garcia-Bustos and Tomasz, 1990; Smith and Klugman, 2001).

An experiment was designed to test whether the order of allele uptake in D39V recombinant strains was crucial for AMX resistance development, and if PBP and MurM substitutions alone were sufficient to explain AMX MICs. Alleles of *pbp2x* and *pbp2b* from AMR37 (*pbp2x37*, *pbp2b37*) and AMR38 (*pbp2x38*, *pbp2b38*) were transformed into D39V and selected on AMX. Strains carrying either a recombinant *pbp2x* or *pbp2b* allele had E-test MICs up to 0.064 µg/mL. However, when both *pbp* alleles were integrated (D39V^2x38-2b38^), the E-test result equalled that of the recombinant allele donor (Figure 7), suggesting that mutations outside these two loci were not involved in the observed reduction in AMX susceptibility in the first-generation strains.

**Figure 7:**
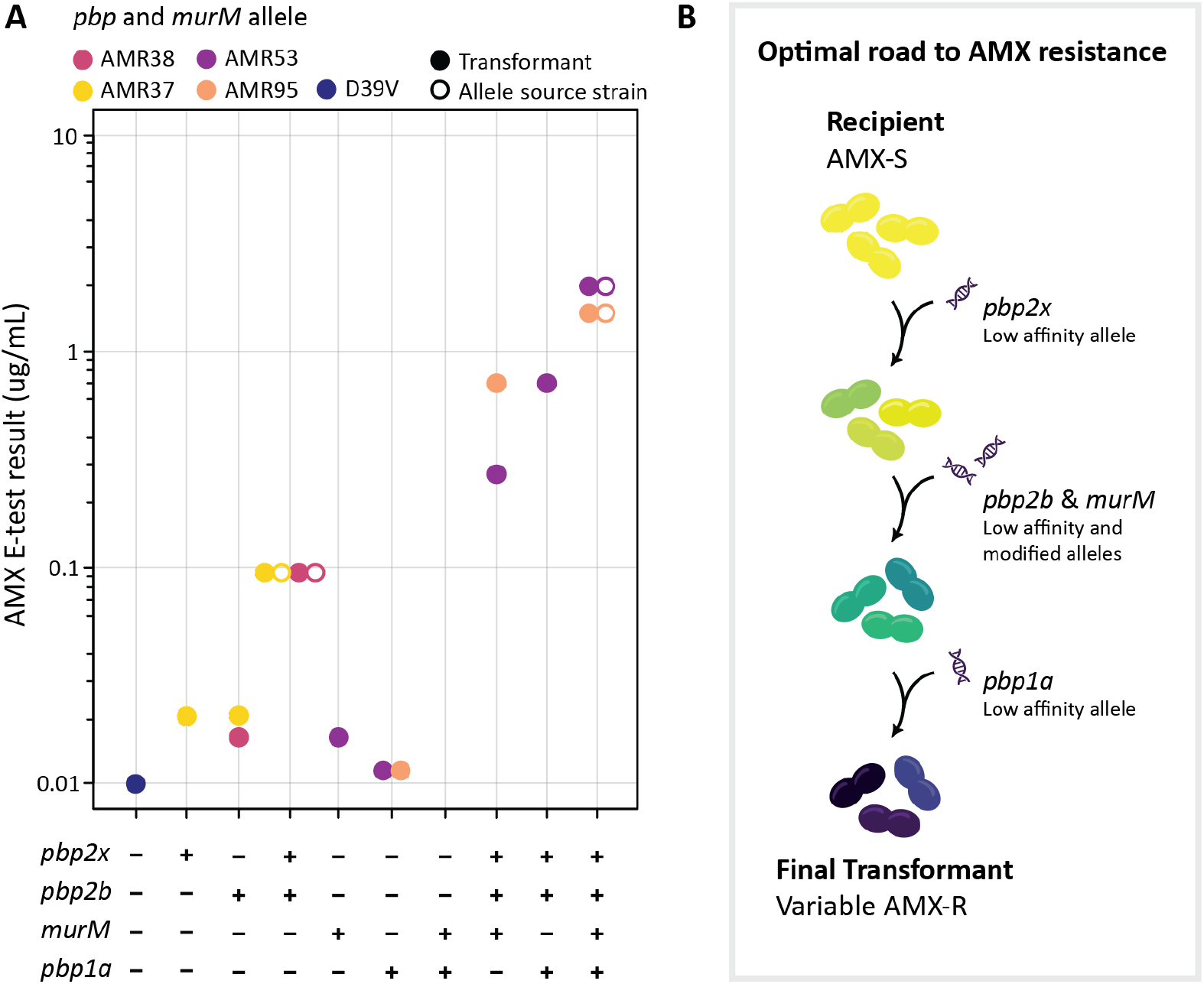
Uptake of mosaic *pbp1a*, *pbp2b*, *pbp2x* and *murM* is required and sufficient for clinically relevant amoxicillin resistance in *S. pneumoniae*. (A) AMX MICs determined by E-test for D39V transformant strains (solid circles) where *pbp* and *murM* alleles from different source strains (empty circles) were transformed in combination. Color shows allele source. (B) Schematic overview of the optimal road to AMX resistance development through horizontal gene transfer. Low affinity *pbp2x* was uptaken in the first round of transformation in both datasets, whereas donor *murM* and *pbp2b* alleles were acquired in the first or second round, depending on recipient and lineage. Low affinity *pbp1a* was recombined in the second round, and only found in strains with AMX MICs above 0.75 µg/mL. In principle uptake of low affinity and modified alleles could be achieved in a single (unlikely) or double step.

To assess whether mutated *pbp2x* and *pbp2b* were critical for *murM* mutation uptake, D39V was transformed with *murM* from both AMR53 (*murM53*) and AMR95 (*murM95*) and MICs determined. Surprisingly, we were able to select for colonies with integration of the AMR53 *murM* allele (D39V^M53^) resulting in an E-test result of 0.023 µg/mL, despite this protein not being a direct target of AMX (Figure 7 and Figure 8A.B). The genome was sequenced, and no spontaneous point mutations were identified in previously described resistance determinants outside *murM* (Table S8). When transformed into the D39V^2x38-2b38^, the E-test MICs reached 0.5 µg/mL and 0.38 µg/mL for AMR95 and AMR53 *murM* alleles (D39V^2x38-2b38-M95^, D39V^2x38-2b38-M53^), respectively. AMX susceptibility decreased substantially when all three genes were mutated, however failed to reach the final MIC of the recombinant allele donor, indicating the necessity of another resistance determinant (Figure 7).

**Figure 8:**
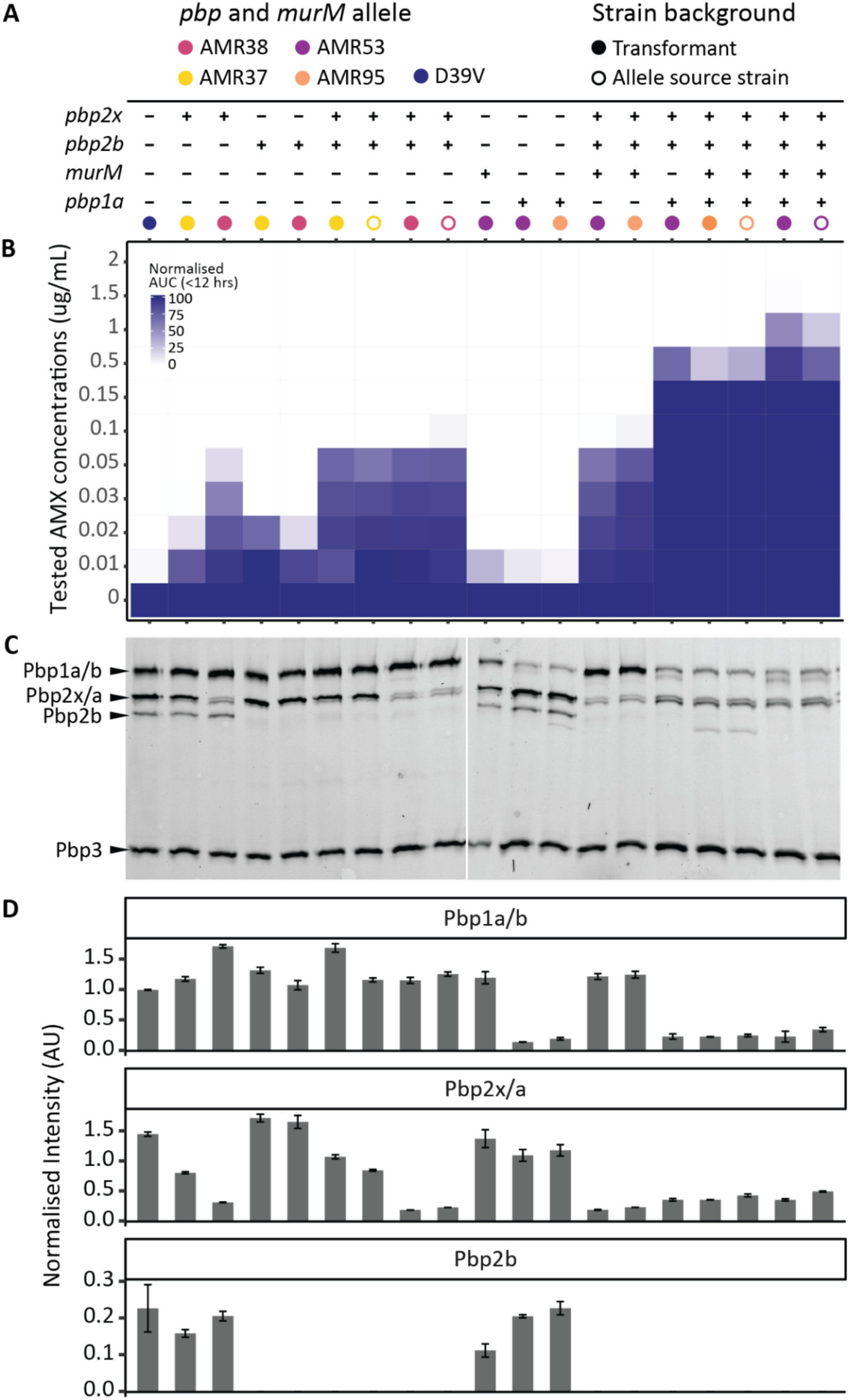
Recombinant PBP proteins visualized by PBP-bocillin binding affinities. Transformed AMX resistant strains produce mosaic PBPs with reduced affinity for Bocillin-FL. Each column shows the phenotypic characterization of one *pbp* and *murM* allele swap strain. **(A)** Table showing the presence (+) and absence (-) of alleles from different source strains (colour). Allele source strains are also included and are indicated by the unfilled circles. **(B)** Survival in different AMX concentrations quantified by determining the area under the growth curve (AUC) after 12 hours growth in AMX normalized by AUC in the absence of AMX. **(C)** Bocillin-FL binding affinity gel. **(D)** Intensity of bands corresponding to Pbp1a/b, Pbp2x/a, and Pbp2b, normalized by the intensity of the Pbp3 band of the same lane, as this allele remained unmodified in all strains.

*pbp1a* alleles from AMR53 and AMR95 (*pbp1a53*, *pbp1a95*) could not be successfully selected on AMX when transformed into D39V, either alone or in conjunction with a *murM* allele from the same donor. Indeed, Pbp1a is not a key target of AMX, and this result was not unexpected. We instead transformed a construct where a kanamycin resistance cassette was cloned behind *pbp1a*, with 1 Kb of homology up- and downstream. Growth in liquid culture of the resulting strains at AMX 0.01 µg/mL was similar to D39V (Figure 8A.B). Interestingly, successful integration occurred for both *pbp1a53* and *pbp1a95* when transformed into D39V^2x38-2b38^ (D39V^2x38-2b38-1a53^, D39V^2x38-2b38-1a95^), suggesting that a mutated *murM* allele was not required for mutations in *pbp1a* to cause a selectable increase in AMX MIC. However, substitutions in Pbp2x, Pbp2b, and Pbp1a alone were insufficient to replicate the E-test results of the allele source strains (Figure 7).

We therefore transformed *murM* and *pbp1a* alleles from AMR53 and AMR95 into D39V^2x38-2b38^ simultaneously, and found that selection was possible on higher concentrations of AMX (1 µg/mL) than for either gene alone (0.2 µg/mL). The resulting strains (D39V^2x38-2b38-M53-1a53^, D39V^2x38-2b38-M95-1a95^) had AMX resistance phenotypes matching those of AMR53 and AMR95, with E-test results of 2 µg/mL and 1.5 µg/mL respectively (Figure 7).

The strains containing all four genes from AMR53 and AMR95 were whole genome sequenced to check for spontaneous point mutations outside these loci. One non-synonymous mutation in *glnA* (glutamine synthase type I) was found in both strains. Downregulation of *glnA* has been linked to the pneumococcal penicillin stress response (el Khoury et al., 2017), but no differences in AMX survival were observed in these strains compared to the allele donors lacking the SNP.

To quantify the affinities of resistant PBP alleles to β-lactam antibiotics, Bocillin-FL binding patterns were determined for allele swap strains and the source strains (Figure 8C.D). This confirmed that the recombinant PBPs had reduced penicillin binding affinities. For Pbp2b, both the AMR37 and AMR38 alleles resulted in almost complete loss of binding, and a band could not be quantified (Figure 8D). A large reduction in affinity was observed for Pbp2x38, larger than for Pbp2x37, which correlated to a small difference in growth at 0.1 µg/mL AMX between the two source strains (Figure 8A.B). This could be explained by 13 amino acid substitutions including S389L and T338A which were not present in the AMR37 allele but have previously been linked to reduced susceptibility to β-lactams (Kocaoglu et al., 2015; Kosowska et al., 2004).

Perhaps most interesting were the two low affinity Pbp1a alleles tested. In all strains containing *pbp1a95* (AMR95 derived), reduced band intensity was observed in the Pbp1a/Pbp1b size range, however, a faster migrating species was also detected. The *pbp1a95* allele contained a spontaneous point mutation (not present in the 11A donor) causing a premature stop codon. This resulted in the loss of 65 amino acids (of a total of 720) and put the predicted size at 73.1 kDa instead of 79.6 kDa. Reduced intensity at the expected Pbp1a/Pbp1b size range was also observed for the AMR53 allele (Pbp1a53) however, an additional Bocillin-FL-tagged species which migrated slightly faster than expected was also observed. The predicted protein length of Pbp1a53 is identical to D39V and the donor, with no premature stop codons identified. The additional band may indicate some degradation of this protein species, although the TP domain must still be available for Bocillin-FL binding. Together, these experiments demonstrate that transfer of the *pbp2X*, *pbp2B*, *pbp1a* and *murM* alleles of 11A into strain D39V is sufficient to explain the observed AMX resistance in transformants obtained using chromosomal DNA.

### High AMX resistance in the D39V comes at a significant fitness loss

Of the 16 recombinant strains sequenced, only EB31 and EB28 from the TIGR4 collection had the same AMX MIC as the donor strain, and a third round of transformation with gDNA into AMR53 failed to increase the MIC. The only region of mutations that EB28 and EB31 had in common that was not present in the highest MIC AMR strains, was a block of 10 mutations in the 590-641 region of Pbp2b (Figure 9C, Table S5). To test whether these substitutions were sufficient to explain the MIC difference, the *pbp2b* allele from EB31 (*pbp2b31*) was transformed into AMR53 and AMR95. The AMX E-test scores for the resulting strains were both 4-6 µg/mL. However, growth curves revealed a significant fitness loss, with a mean maximum specific growth rate of 0.11 µ/h for AMR95 decreasing almost 100-fold to 0.0036 µ/h for AMR95^2b31^, which was exacerbated by the presence of AMX (Figure 9B). In contrast, AMR53^2b31^ grew almost 3-fold slower than AMR53 (AMR53 0.12 µ/h, AMR53^2b31^ 0.046 µ/h, Figure 9B), but showed a lytic phenotype in stationary phase. Interestingly, this strain outgrew the 11A donor in sublethal AMX concentrations (Figure 9A). The difference in growth rate between AMR95^2b31^ and AMR53^2b31^ was likely explained by the truncated Pbp1a95 protein. Significant modifications to the active sites of essential proteins often need to be compensated for, which may be impacted by the sub-optimal performance of other proteins in the system. Although the TP and TG domains of Pbp1a95 were intact, it is possible that the loss of 65 amino acids from the C-terminal resulted in an overall reduction in function, unrelated to its AMX binding affinity. This correlates with previous studies showing that compensation by substitutions in Pbp1A are required to mitigate fitness loss resulting from modifications to the Pbp2b active site (Albarracín Orio et al., 2011). Together, these experiments show that D39V is capable of reaching high AMX resistance, but that the absence of compensation by Pbp1A substitutions may result in significant fitness costs.

**Figure 9:**
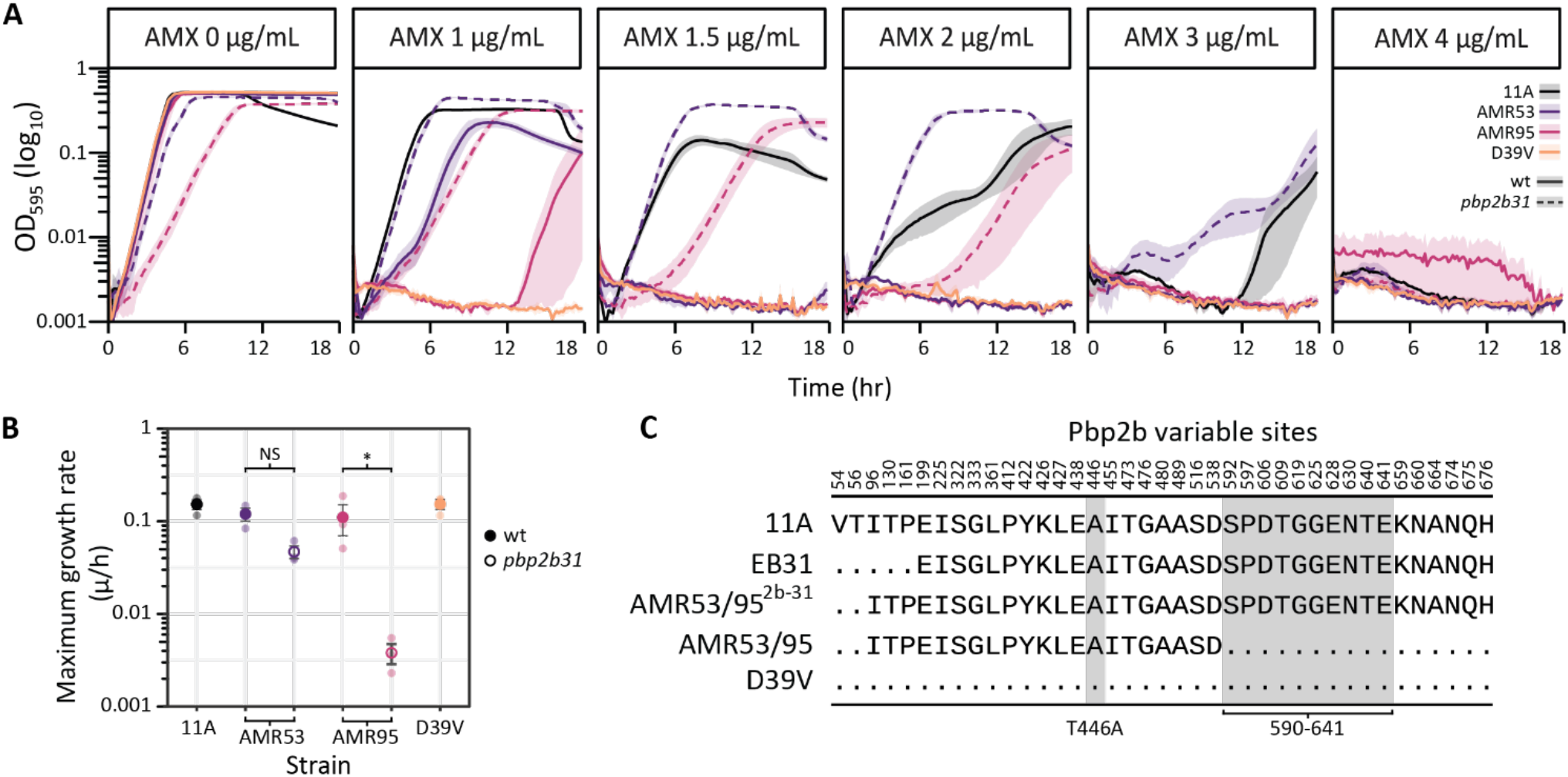
Ten substitutions in Pbp2b 590-641 region essential for high AMX resistance. **(A)** Growth of AMR53^2b31^ and AMR95^2b31^ (dotted lines) in different AMX concentrations. 11A, D39V, AMR53, and AMR95 (solid lines) are shown as controls. **(B)** Dot plot of maximum specific growth rates (µ/h) calculated from the AMX 0 µg/mL condition in triplicate (transparent circles). The mean for each strain is shown by an opaque circle, solid for wild-type (wt) strains and empty for AMR53^2b31^ and AMR95^2b31^. Color denotes strain background. Significance was determined by one-way ANOVA with a Tukey correction for multiple testing. **(C)** Amino acid sites in pbp2b which differ from D39V for strains 11A, EB31, AMR53 (identical to AMR95), and AMR53^2b31^ (the result of transforming AMR53 or AMR95 with the EB31 pbp2b allele). Dots indicate the amino acid matches D39V, and key substitutions, T446A and a block of 10 between 590-641 (Cafini et al., 2006; du Plessis et al., 2002; Kosowska et al., 2004), are highlighted in grey.

## Discussion

Natural competence and homologous recombination in *S. pneumoniae* allow for enormous genomic plasticity which contributes to the rapid development of vaccine-escape and antibiotic resistant isolates. In one round of transformation, more than 29 recombination events can occur in a single cell (Cowley et al., 2018), while events up to 50 Kb in length have been identified in clinical isolates (Wyres et al., 2013), allowing for significant remodelling at multiple loci simultaneously. Indeed, here we show a two-step sequential transformation experiment with up to 20 individual recombination events around the genome and lengths up to 13 Kb, resulting in an average donor genome transfer per step of 0.38%. This allows for huge genomic diversity, which, when placed under selective pressure, can lead to the rapid spread of antibiotic resistance-associated genotypes (Chewapreecha et al., 2014).

β-lactam resistance in the pneumococcus relies on DNA movement between closely related strains and species within the same niche, allowing tens of substitutions to accumulate in the target proteins (Hakenbeck et al., 2012). It has previously been shown through *in vitro* transformation experiments into a laboratory strain that the main determinants of β-lactam resistance are modified alleles of *pbp2x*, *pbp2b*, *pbp1a*, and *murM* (Fani et al., 2014; Smith and Klugman, 2001; Trzciński et al., 2004). For the front-line antibiotic AMX, resistance has also been previously associated with mutations in these alleles although the relative importance of the changes in each locus is not well understood (Cafini et al., 2006; Chesnel et al., 2005; du Plessis et al., 2002; Garcia-Bustos and Tomasz, 1990; Kosowska et al., 2004; Smith and Klugman, 2001). We confirmed the requirement for mutations in all four genes and showed that subtle differences between alleles could cause noticeable changes in both AMX-dependent fitness and the penicillin-binding affinity of PBPs.

In addition, a specific order of mutation uptake for penicillin resistance has been demonstrated in both *Staphylococcus aureus* and the pneumococcus (Demerec, 1945; Hotchkiss, 1951; Shockley and Hotchkiss, 1970), and for AMX this order was found to be *pbp2x*, *pbp2b*, then *pb1a*, with the role of *murM* unable to be confirmed in that experimental system (du Plessis et al., 2002). Here, we confirm that substitutions in Pbp2x are a critical first step for AMX resistance, perhaps reflecting the protein’s high affinity for the antibiotic (Kocaoglu et al., 2015)(Figure 7C), and that the T338A substitution is required for resistance to fully develop. We also show that mutations in both *pbp2b* and *murM* are essential intermediary steps, and that a modified *pbp1a* allele is crucial for high-level AMX resistance. It is however important to note that these experiments were performed using high concentrations of naked DNA as a model, while in the nasopharynx genetic exchange likely occurs via fratricide or from small quantities of free environmental DNA (Eldholm et al., 2009; Johnsborg et al., 2008). It may be that lower donor DNA concentrations cause slower resistance allele acquisition, and result in lower fitness costs. In addition, previous studies have strongly implied the existence of multiple mechanisms of AMX resistance in *S. pneumoniae*, associated with different *pbp2b* and *murM* alleles (Cafini et al., 2006; Càmara et al., 2018; du Plessis et al., 2002), and that by using a single donor strain we were limited to the study of only one. However, we confirmed that a previously described block of ten substitutions in the 590-641 region of the Pbp2b TP domain (Cafini et al., 2006; du Plessis et al., 2002; Kosowska et al., 2004) was essential to reconstruct the AMX MIC of the serotype 11A ST6521 donor strain in D39V and TIGR4. Perhaps most interestingly, we also show that the loss of 65 amino acids from the C-terminus of Pbp1a results in a significant fitness cost in the presence of these additional Pbp2b substitutions, highlighting the importance of interactions and fitness loss compensation between essential cell-wall synthesis proteins (Albarracín Orio et al., 2011).

In addition to these definite experiments demonstrating the evolutionary route to clinical AMX resistance, we show that non-contiguous recombination could result from either a single, or multiple, donor DNA molecules. It is easy to imagine a long ssDNA-RecA presynaptic filament interacting with the chromosome at multiple points in the 3-dimensional cellular space (Forget and Kowalczykowski, 2012; Yang et al., 2020)(Figure 1 model), which then initiates the formation of several D-loops simultaneously as homologous DNA-DNA interactions stabilize (Bell and Kowalczykowski, 2016; Hsieh et al., 1992; Mazin and Kowalczykowski, 1996). On the other hand, SsbB coated ssDNA sequestered in the cytoplasm could provide a pool from which multiple donor molecules homologous for the same chromosomal region could be sourced (Attaiech et al., 2011). However, we also observed that both transformation efficiency and non-contiguous event occurrence increased with length of homology and total length of the donor molecule, suggesting that single donor molecule events may simply be more likely. This concurs with the previous observation of transforming DNA concentration-independent differences in recombination event density, which would not have been expected under a multiple donor molecule hypothesis (Croucher et al., 2012). Recombination structures such as these have also been observed in *H. influenzae*, with the authors noting that localized clustering of long tracts of donor sequence appeared too close to have occurred by chance (Mell et al., 2011). They hypothesized incoming DNA degradation via cytosolic or translocation endonucleases resulted in these clusters. In addition, mismatch repair via the Hex system could in principle be responsible for the reversion of small numbers of point mutations between non-contiguous recombination segments, although the system was likely saturated in our model (Humbert et al., 1995). To untangle the different possible mechanisms of non-contiguous recombination from other effects on transformation efficiency, testing mixtures of different donor fragment lengths would be necessary.

Altering the active sites of essential proteins in cell division is risky and requires a delicate balance between antibiotic avoidance and fitness loss. Accumulation of sufficient mutations in the appropriate order is aided both by the simultaneous recombination of multiple donor fragments distally around the genome, and by the occurrence of complex non-contiguous events. The experimental transformation experiments combined with whole genome sequencing as performed here provide valuable insights into viable and non-viable evolutionary paths toward AMX resistance development in the pneumococcus. The strong co-dependence among different proteins and their alleles creates significant challenges for complete understanding of resistance mechanisms, and emphasizes the importance of combining data from phenotypic, molecular, genomic, and genome-wide association studies.

## Methods

### Bacterial strains, growth conditions, and antibiotics

*S. pneumoniae* strains are shown in Table 1 and Table S1. Bacteria were cultured in liquid C+Y medium with no shaking at 37°C. C+Y media was adapted from Adams and Roe (Martin et al., 1995).

For transformation, *S. pneumoniae* was grown in C+Y (pH 6.8) at 37°C to OD_595_ 0.1. Competence was induced by addition of 100 ng/ml synthetic CSP-1 (D39V) or CSP-2 (TIGR4) and 12 min incubation at 37°C. DNA uptake occurred during 20 min at 30°C, followed by dilution and recovery for 1.5 hr. Transformants were selected by plating inside Columbia agar supplemented with 3% defibrinated sheep blood (CBA, Thermo Scientific) containing the appropriate antibiotic and incubated at 37°C in 5% CO_2_ overnight. Successful transformants were confirmed by PCR and Sanger sequencing (Microsynth).

Strains were stocked at OD_595_ 0.3 in C + Y with 15% glycerol at −80°C.

AMX (Sigma Aldrich) powder was dissolved initially in 100% DMSO. This solution was diluted in molecular grade water with 4% DMSO to two concentrations, 100 µg/mL and 1 mg/mL, and stored at −80°C.

### Minimum Inhibitory Concentrations (MICs)

Bacteria were grown in C+Y broth to OD_595_ 0.1, then AMX MIC was determined using E-tests (Biomeriaux, Lyon) on Mueller Hinton agar containing 5% defibrinated sheep’s blood. MICs were read after 18 h incubation at 37°C and 5% CO_2_.

Strains with MICs below the limit of the E-tests (0.016 µg/mL) were tested in Mueller Hinton cation-adjusted broth with 5% defibrinated sheep’s blood using the broth micro-dilution method (Charles et al., 2016). Briefly, bacteria were streaked onto MH2-blood plates and grown overnight. Colonies were picked from plates and ressupended in PBS (pH 7.4) to McFarlane standard 0.5. This inoculum was diluted 100-fold into 96-well plates containing MH2-blood broth and AMX (in 2-fold concentration steps) then plated on agar plates as a control for contamination and cell density. Plates were incubated for 18 hours at 37°C and 5% CO_2_. MIC was defined as the lowest concentration of antibiotic where visible α-haemolysis occupied approximately less than 10% of the surface area of the well. Due to a high level of subjectivity, MIC determination was repeated twice on different days, and MIC calls were confirmed by a second person in each case.

### Growth curves and analysis

Strains were precultured in C+Y broth to OD 0.1, then diluted 100-fold into 96-well microtiter plates, containing fresh C+Y and AMX. The OD_595_ was measured every 10 min for 12 hr in a plate reader (Infinite F200, Tecan) with incubation at 37°C. All strains were tested in triplicate Area under the curve was determined using TecanExtracter (Dénéréaz, unpublished). The AUC of each replicate was divided by that of the no antibiotic control for each strain, to normalize for strain-dependent differences in growth. The mean and standard error of the mean (SEM) of the normalized AUC was determined for three biological replicates.

Statistical significance of maximum specific growth rate (µ/h) was determined using one-way ANOVA with a Tukey correction for multiple testing.

### Bocillin-FL binding assay

Bocillin-FL is a fluorescently labelled penicillin molecule which can be used to visualise and quantify PBP-penicillin binding affinity *in vitro*. Strains were grown to OD 0.2 in 4 mL C+Y, washed and resuspended in phosphate buffered saline (PBS), then incubated for 30 min at 37C in 5 µg/mL bocillin-FL (Invitrogen). Cells were washed in ice cold PBS, then lysed by sonication (10 x 1s pulses at 30% power). The cell lysate was spun down and supernatant removed to collect the membrane fraction, which was homogenized by sonication (1s pulse at 10%). Total protein concentration was determined by nanodrop, then samples were diluted in NuPAGE LDS sample buffer (Life Technologies) with 10 mM DTT and incubated at 95°C for 5 min. Samples were loaded into two 15-well Novex™ WedgeWell™ 10% Tris-Glycine 1.0 mm Mini Protein Gels (Invitrogen), to a total protein amount of approximately 1.8 mg in a maximum volume of 30 µL. Where necessary, gels were imaged simultaneously on an Amersham Typhoon (GE Healthcare) with Cy2 DIGE filter setup. Bands were quantified using ImageQuant (GE Healthcare). As all strains tested had identical PBP3 proteins, the intensities of the PBP1a/b, PBP2x/a, and PBP2b bands were divided by the PBP3 band intensity of the corresponding sample for normalisation. Imaging and quantification were performed in duplicate. Coomassie Blue staining was also performed, to confirm equal sample loading.

### Phase-contrast time-lapse microscopy

For time-lapse microscopy, cells were pre-cultured in liquid C + Y medium at 37°C and spotted on a low-melting 1.2% C + Y agarose patch, with or without AMX at 1 µg/mL. Phase-contrast pictures were taken every 10 min as described (de Jong et al., 2011) using a Leica DMi8 with a 100x phase contrast objective.

### Isolation of genomic DNA

10 mL of OD_595_ 0.2 culture was pelleted by centrifugation, then resuspended in Nuclei Lysis solution (Promega) supplemented with 0.05% SDS, 0.025% deoxycholate (DOC), and 200 µg/mL RNase A. This was incubated at 37°C for 20 minutes, 80°C for 5 minutes, then 37°C for 10 minutes. Protein Precipitation Solution (Promega) was added to the lysate, vortexed vigorously, then incubated on ice for 10 minutes. The precipitated protein was pelleted by centrifugation and the supernatant transferred into isopropanol to precipitate the DNA, which was then collected by centrifugation. DNA was washed once in 70% ethanol before air drying and resuspension in the appropriate buffer (either TE buffer or molecular grade water). DNA was run on a 1% agarose gel to confirm no significant degradation had occurred, and stored at 4°C.

### Serial Transformation of S. pneumoniae with genomic DNA

AMX susceptible recipient strains were transformed with 100 μg/mL genomic DNA isolated from AMX resistant strain 11A (Table 1) as above, with the following modifications. DNA uptake occurred during 30 min, followed by recovery for 2 hrs. Transformants were selected by plating inside Columbia agar supplemented with 5% defibrinated sheep blood and a range of AMX (0.03 µg/mL). Ten colonies were then picked at random and streaked onto CBA plates containing AMX. The next day, single colonies were grown up in C+Y with AMX and stocked. MICs were determined for these 10 strains, which were then subjected to another round of transformation and selection on a range of AMX concentrations (0.17 - 0.5 µg/mL), and 5-10 single colonies were picked for each of the 10 strains transformed. These were isolated, and MICs determined, as above. To test for significant differences between MICs of second-generation strains in different lineages, a Fisher’s Exact Test with Monte Carlo p-value simulation was used.

To control for spontaneous mutations arising during transformation and recombination, cells transformed with either no DNA, or with D39V genomic DNA, were also plated in CBA with AMX (0.03). Neither resulted in visible colonies.

### Whole genome sequencing and quality control

Genomic DNA from recombinant strains from each recipient background, as well as 11A and TIGR4 were Illumina sequenced (PE150, Novogene and Eurofins Genomics), with a mean coverage of 600-fold. Reads were trimmed using Trimmomatic (Bolger et al., 2014), then assembled using SPAdes (Nurk et al., 2013). Assembly quality was assessed with Quast (Gurevich et al., 2013), and read depth was determined in samtools (Li et al., 2009). Previously published D39V Illumina reads were filtered as above and used where necessary (GEO accessions GSE54199 and GSE69729) (Slager et al., 2014). Allelic exchange strains (D39V^M53^, D39V^2x38-2b38-M53-1a53^, D39V^1a53^, D39V^2x38-2b38-M95-1a95^) were treated in the same way, then variants were called using bbmap (Bushnell, 2013), Freebayes (Garrison and Marth, 2012), and vcftools (Danecek et al., 2011), in order to check for random mutations arising elsewhere in the genome.

11A, TIGR4, and a subset of recombinant strains were sent for PacBio sequencing (Sequel I and II, Lausanne Genomic Technologies Facility). Demultiplexing and quality control were performed using SMRTlink (PacBio), then reads shorter than 1000 bp were removed with filtlong (Wick, 2017). Recombinant strain reads were mapped to the recipient reference genome using NGMLR and structural variants (SVs) called with sniffles with default settings, except with read mapping quality > 40 and read depth > 50 (Sedlazeck et al., 2018). Imprecise SVs were then filtered out. DNA methylation detection was performed using motifMaker (Clark et al., 2012).

Raw read files are deposited with NCBI Sequence Read Archive under BioProject PRJNA789167 (Individual SRA accession numbers for recombinant strains can be found in Table S4).

### Hybrid assemblies of reference genomes

As PacBio sequencing depth was greater than 200x, genomes were assembled in a long-read only manner using Trycycler (Wick et al., 2021), followed by short-read polishing.

Briefly, filtered reads were subsampled 12 times and *de novo* assembled in four different assemblers, flye (Kolmogorov et al., 2019), raven (Wang et al., 2018), wtdbg2 (Ruan and Li, 2020), and miniasm plus minipolish (Li, 2018, 2016). Some assemblies were removed during the Trycycler cluster and reconcile steps, leaving the final consensus sequence to be built from seven to eight assemblies. The consensus assembly was subjected to rounds of polishing with PacBio reads using pbmm2 and pbgcpp (PacBio Toolkit), followed by one round of short-read polishing with Pilon (Walker et al., 2014).

Genomes were annotated in Prokka (Seemann, 2014) using Barrnap and Aragorn (Laslett and Canback, 2004; Seemann, 2019) and are deposited at NCBI under BioProject PRJNA789167 (accession numbers currently in process).

### Recombination detection with NGS

Competitive mapping of filtered illumina reads onto a pseudogenome containing both recipient and donor chromosomes was used for crude visualization of recombination loci (bbmap) (Matthey et al., 2019). In order to characterize these loci more accurately, filtered reads of recombinant and recipient strains were mapped onto the 11A hybrid genome assembly using smalt, then variants were called with bcftools (Li, 2011). SNPs were filtered as described previously (Cowley et al. 2018), and used to build a consensus alignment of donor, recipient, and recombinant strains. The consensus alignment was run through a python script designed to identify SNPs between recombinant and donor which are not shared with the recipient genome. Once identified, the match was extended until the donor and recombinant sequences are no longer identical, then the start, end, and length of detected recombination events are extracted (Cowley et al 2018).

### Identifying non-contiguous recombination events

Many recombination events were located very close together, with sometimes only a few recipient allele SNPs separating the two, suggesting a single donor DNA molecule as the source. To determine if events were significantly close together, a bootstrapping approach was applied (Croucher et al., 2012). Firstly, the distances between one recombination event and each of the other recombination events in a single genome were calculated. The circular chromosome gives two values, and the shortest distance between any two events was added to the test population. This was calculated for every sequenced strain in the D39V and TIGR4 recombinants, and for second-generation strains any prior events carried through from the first-generation were removed from the dataset. The shortest recombination distance within a strain became the test value (*d_test_*), and then the test population was sampled (with replacement).

The distribution of a bootstrapped sample (same size as the test population) was used to test the hypothesis that *d_test_* was significantly shorter than expected under the null hypothesis that recombination events are located at random around the chromosome. A Holm correction was applied to account for multiple testing, based on the number of recombination events in the strain. Bootstrap tests were performed one thousand times for each *d_test_* value and considered significantly short if the null hypothesis was rejected at least 95% of the time. If a *d_test_* value was significant, then the distance between the next shortest pairing within the strain became *d_test_*, and was tested in the same fashion. This continued until a *d_test_* where the null hypothesis could not be rejected.

When recombination events were considered significantly close together, they were linked together into non-contiguous recombination events.

### Confirming the origins of non-contiguous recombination events in vivo

A 4.25 Kb region of the 11A genome spanning *pbp2x* was used to test the origin of non-contiguous recombination. This region included an 846 bp stretch with SNPs conferring a small decrease in AMX susceptibility sufficient for selection. One long fragment spanning the entire region was amplified with OVL5375 and OVL5378. Two shorter fragments were then amplified, an upstream fragment (Up) with OVL5379 and OVL5375 (2.84 Kb) and a downstream fragment (Down) with OVL5376 and OVL5378 (2.26 Kb), which overlapped at the SNPs determined to be necessary for selection (Table S2, S5). The three donor DNA templates used in the experiment and their SNPs are shown in Figure 6C (Table S6).

D39V was transformed as above, with equimolar quantities of each fragment separately, as well as a combination of the two short fragments, and a negative control with no DNA. Transformants were selected on AMX (0.015 µg/mL), and the transformation efficiency was determined by dividing by the total viable count. Individual colonies were restreaked on AMX, then single colonies were picked and the 4.25 Kb region of interest was amplified and Sanger sequenced (Microsynth, Table S2 oligos). Nucleotide sequences were aligned in Mega (Kumar et al., 2018).

As molecule length affects transformation efficiency (Keller et al., 2019; Kurushima et al., 2020; Lee et al., 1998), an additional experiment was performed, where the amplified fragments were all extended to 6 Kb by the addition of non-homologous DNA amplified from *E. coli* and ligated to one end using Golden Gate assembly with BsmBI and T4 DNA ligase (NEB, Vazyme). All oligos and the corresponding templates used in the cloning for these experiments are outlined in Table S6, and a comprehensive oligo list can be found in Table S2. The assemblies were then amplified to ensure high quality of transforming fragments, resulting in three fragments of equal size, with differing lengths of homologous regions to the *pbp2x* locus. These were then transformed, amplified, and Sanger sequenced as above, and the transformation efficiency determined.

### Gene and protein alignments

Nucleotide sequences of *pbp* and *murM* genes were translated using Emboss:TransSeq (Madeira et al., 2019). Alignments were performed with Muscle (Edgar, 2004). Phylogenetic trees were produced using concatenated protein alignments of Pbp2x, Pbp2b, Pbp1a, and MurM as input for FastTree (default settings), and visualised in ITOL (Letunic and Bork, 2021; Price et al., 2009).

### Amplification and transformation of *pbp* and *murM* loci for allelic exchange experiments

Genes for *pbps* were amplified with Phanta Max Super-Fidelity DNA Polymerase (Vazyme) and primers ordered from Sigma (Table S2). Equimolar quantities of PCR products were used for transformation. To avoid transforming mutations in *murN* alongside those in *murM*, alleles at this locus were cloned into a construct with D39V flanking sequences. The upstream and downstream fragments were amplified from D39V with OVL5540/OVL5779 and OVL5780/OVL5541 respectively, and the *murM* allele from AMR53 or AMR95 with OVL5777/OVL5778. Fragments were then assembled using BsmBI and T4 DNA ligase. *pbp2x* and *pbp2b* alleles from AMR37 and AMR38 were transformed into D39V as above, selected on 0.012 µg/mL, 0.015 µg/mL, 0.02 µg/mL AMX, and confirmed by Sanger sequencing. D39V^2x38-2b38^ (Table 1) was then used as the recipient for *murM* and *pbp1a* alleles from AMR53 and AMR95, with selection on 0.2 µg/mL, 0.5 µg/mL and 1 µg/mL AMX.

## Acknowledgements

We would like to thank Mark van der Linden at the German National Reference Center for Streptococci for the 11A (SN75752) clinical isolate, and the Lausanne Genomic Technologies Facility for their valuable work in PacBio sequencing and data quality control. We would also like to thank Arnau Domenech Pena, Alice Wallef, Vincent de Bakker, Julien Denereaz, and the rest of the Veening Lab for their help, contributions, and feedback.

P.G. was supported by the University of Lausanne Faculty of Biology and Medicine PhD fellowship. Work in the Veening lab is supported by the Swiss National Science Foundation (SNSF) (project grant 310030_192517), SNSF JPIAMR grant (40AR40_185533), SNSF NCCR ‘AntiResist’ (51NF40_180541) and ERC consolidator grant 771534-PneumoCaTChER.

## Supplementary Tables

**Table S1: Recombinant strains isolated in sequential rounds of transformation with AMX resistant DNA and their associated MICs.**

**Table S2: Primers used in this study.**

**Table S3: List of all recombination events identified in transformants using SNPs.**

**Table S4: Summary statistics for the characterisation of SNP and recombination results for sequenced recombinant strains.** (A) All predicted events detected when compared to parent strain genome. Potential non-contiguity not accounted for. (B) Total Dna transferred during the final round of transformation for that strain. (*) All DNA or SNPs transferred into the original recipient (D39V or TIGR4), through one or two rounds of transformation (depending on the generation). SRA accession numbers for raw Illumina and PacBio reads used for analysis are also provided.

**Table S5: Key amino acid substitutions in PBPs and MurM of recombinant strains.** Substitutions listed have been associated with β-lactam resistance in the literature previously. Cells coloured green have the same allele as D39V, those coloured blue have the same allele as 11A. Strain AMX MICs are also indicated for reference.

**Table S6: Cloning design for fragments used to test for single-molecule non-contiguous recombination at the *pbp2x-mraY* locus.**

**Table S7: Summary statistics from colonies isolated after transformation with different length and combinations of donor fragments to explain the occurrence of non-contiguous recombination.**

**Table S8: SNPs detected in D39V^M53^.**

## Supplementary Figures

**Figure S1: Fluorescence and coomassie stained images of Bocillin-FL gels.** (A) Bocillin-FL gel from Figure 3 and (B) gels from Figure 8.

**Figure S2: Identification of non-contiguous recombination from sequenced recombinant strains using a bootstrapping method** (Croucher et al. 2012). (A) Distances between recombination events which were found to shorter than expected under a null hypothesis where events occur randomly around the chromosome. (B) Schematic showing how events were treated if they were found to be closer to each other than expected. (C) Histogram of recombination events length across all sequenced recombinant strains if non-contiguous recombination is not taken into account. (D) Histogram of recombination events length across all sequenced recombinant strains if non-contiguous recombination is accounted for.

**Figure S3: Alignments of pbp2x-mraY from colonies isolated after transformation with different length and combinations of donor fragments to explain the occurrence of non-contiguous recombination.** Facets show colonies selected from different donor templates which correspond to Figure 6. Bases which match the recipient (D39V) are coloured grey, those which match the donor (11A) are shown in blue. (A) Long, (B) Up, (C) Down, (D) Up & Down, (E) Long-NH, (F) NH-Up, (G) Down-NH, and (H) NH-Up and Down-NH.

**Figure S4: Dendrogram of strain relationships built from amino acid sequences of Pbp and murM proteins.** Alignments of Pbp1a, Pbp2b, Pbp2x, and MurM amino acid sequences were used to construct a phylogenetic tree of donor, recipient, and recombinant strains. AMX MICs are shown on the right, colour scale represents MIC and corresponds to figure 2, where yellow is an AMX MIC of 0.01 µg/mL and dark purple is an MIC of 4 µg/mL.

