## Supplemental Figures S1-S4 for "The acquisition of clinically relevant amoxicillin resistance in *Streptococcus pneumoniae* requires ordered horizontal gene transfer of four loci"

### Supplementary Figures

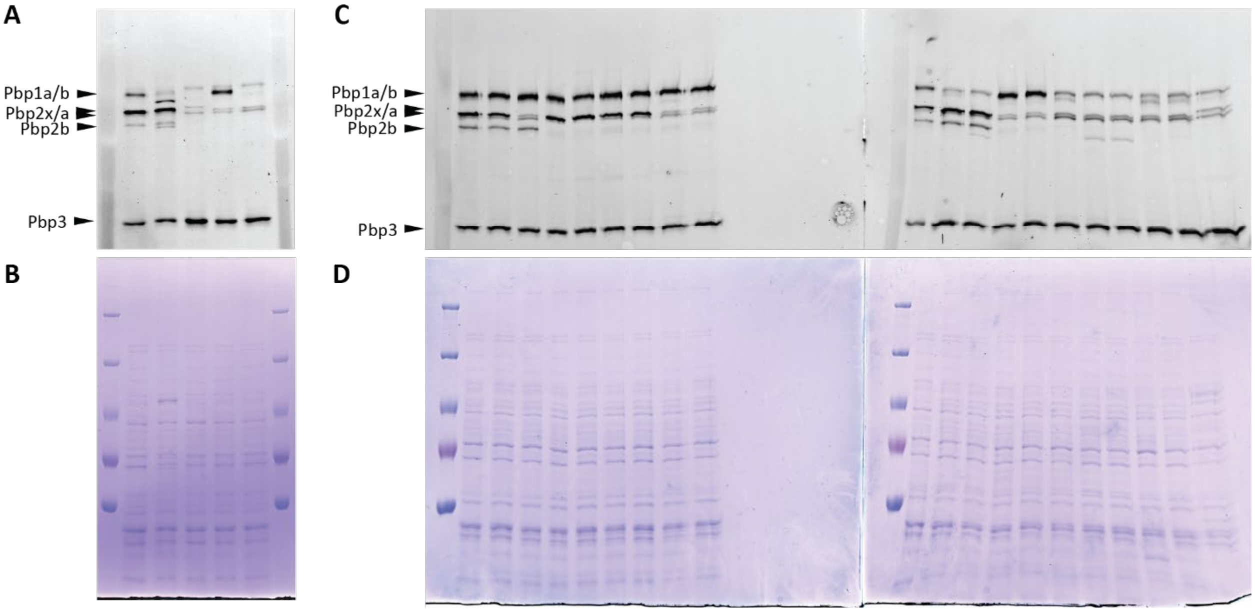


**Figure S1: Fluorescence and coomassie stained images of Bocillin-FL gels**. (A) Bocillin-FL labelled cell extracts imaged in an Amersham Typhoon with Cy2 filter setup showing PBP binding affinities, from Figure 3. (B) Coomassie stain of Bocillin-FL gel from Figure 3. (C) Bocillin-FL labelled cell extracts imaged in an Amersham Typhoon Cy2 filter setup from Figure 8. (D) Coomassie stain of Bocillin-FL gel from Figure 8.

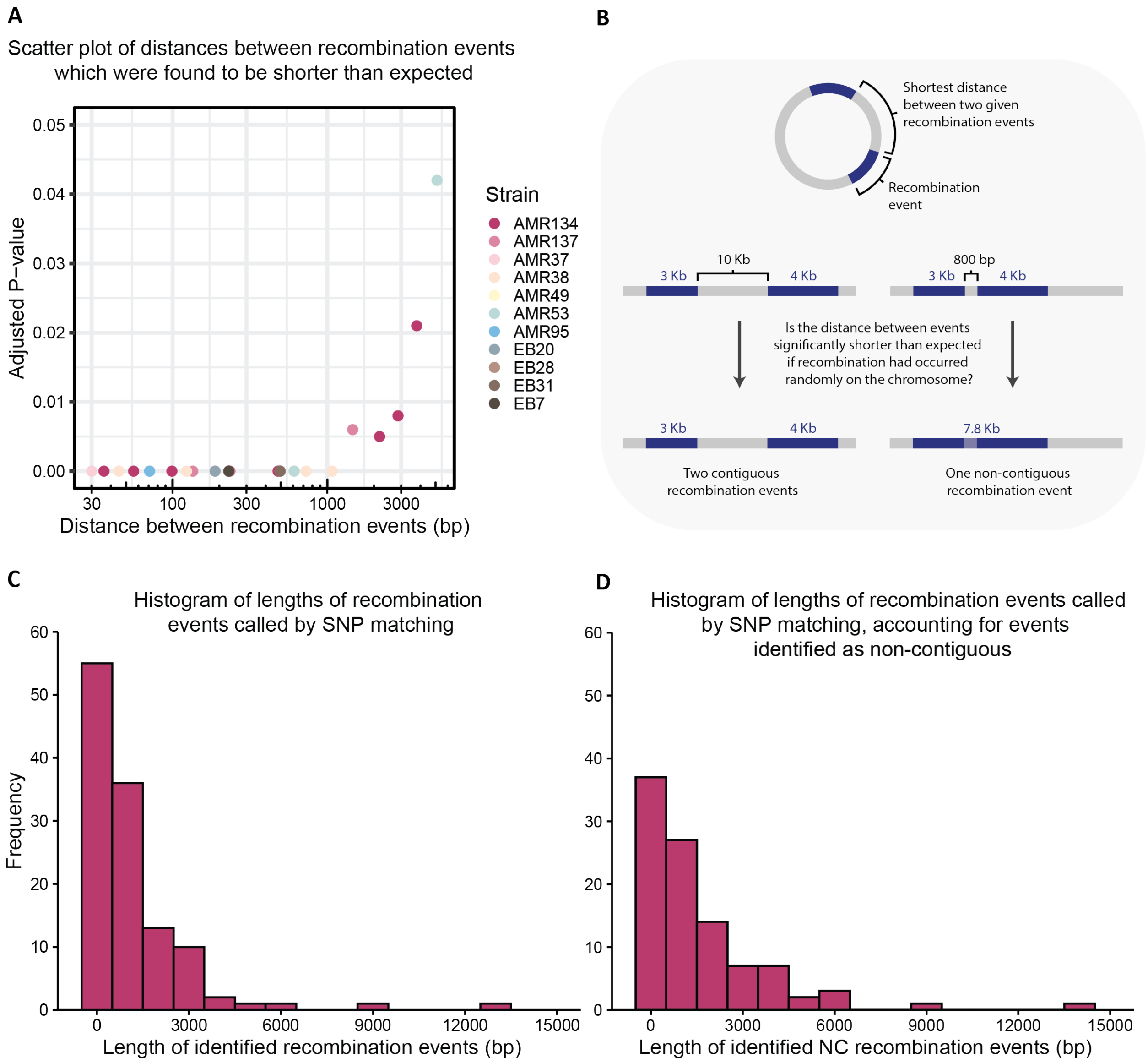


**Figure S2: Identification of non-contiguous recombination from sequenced recombinant strains using a bootstrapping method** (Croucher et al. 2012). (A) Distances between recombination events which were found to shorter than expected under a null hypothesis where events occur randomly around the chromosome. (B) Schematic showing how events were treated if they were found to be closer to each other than expected. (C) Histogram of recombination events length across all sequenced recombinant strains if non-contiguous recombination is not taken into account. (D) Histogram of recombination events length across all sequenced recombinant strains if non-contiguous recombination is accounted for.

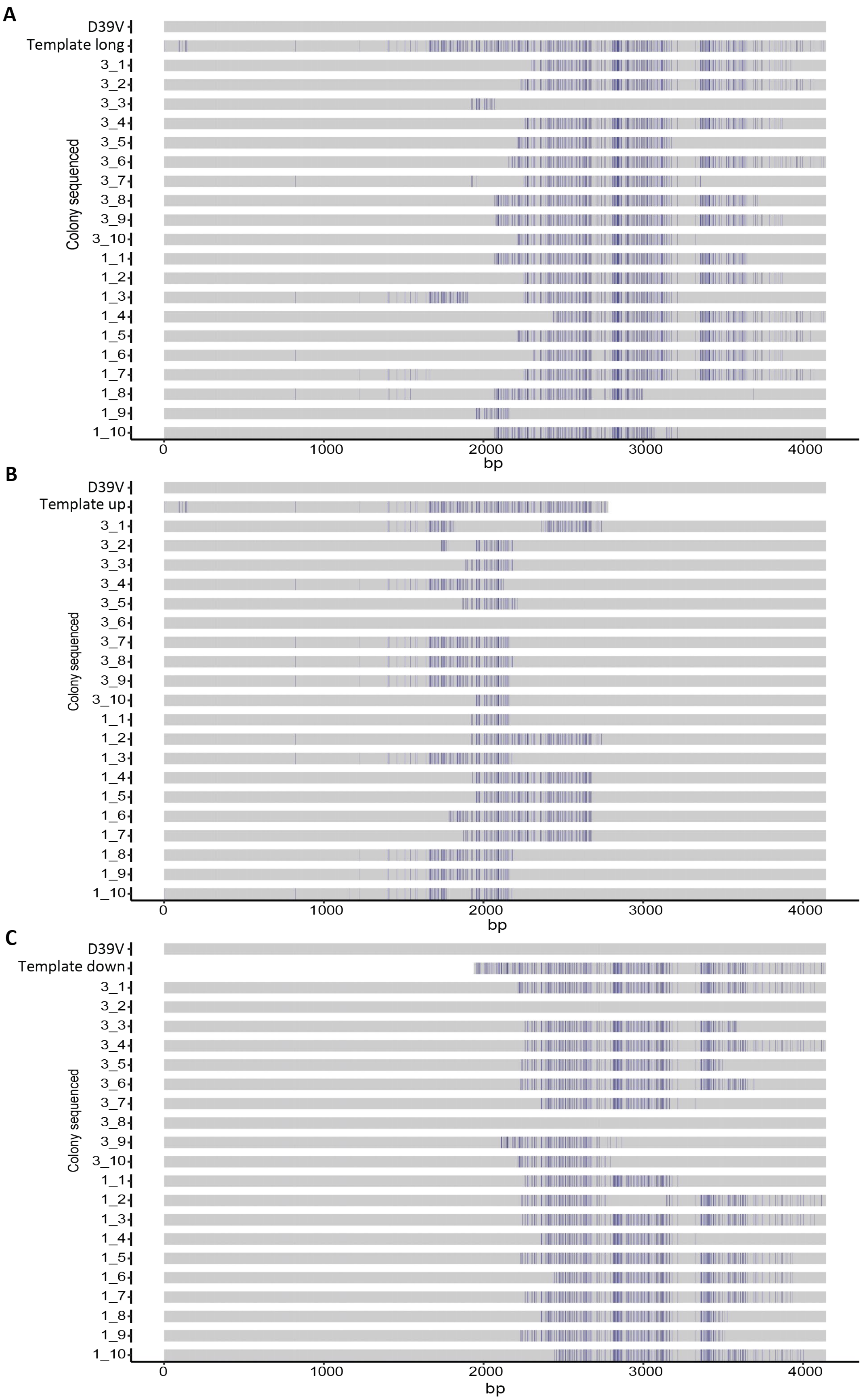

A
D39V
Template long
3_1
3_2
3_3
3_4
3_5
3_6
3_7
3_8
3_9
3_10
1_1
1_2
1_3
1_4
1_5
1_6
1_7
1_8
1_9
1_10
Colony sequenced
0
1000
2000
3000
4000
bp
B
D39V
Template up
3_1
3_2
3_3
3_4
3_5
3_6
3_7
3_8
3_9
3_10
1_1
1_2
1_3
1_4
1_5
1_6
1_7
1_8
1_9
1_10
Colony sequenced
0
1000
2000
3000
4000
bp
C
D39V
Template down
3_1
3_2
3_3
3_4
3_5
3_6
3_7
3_8
3_9
3_10
1_1
1_2
1_3
1_4
1_5
1_6
1_7
1_8
1_9
1_10
Colony sequenced
0
1000
2000
3000
4000
bp

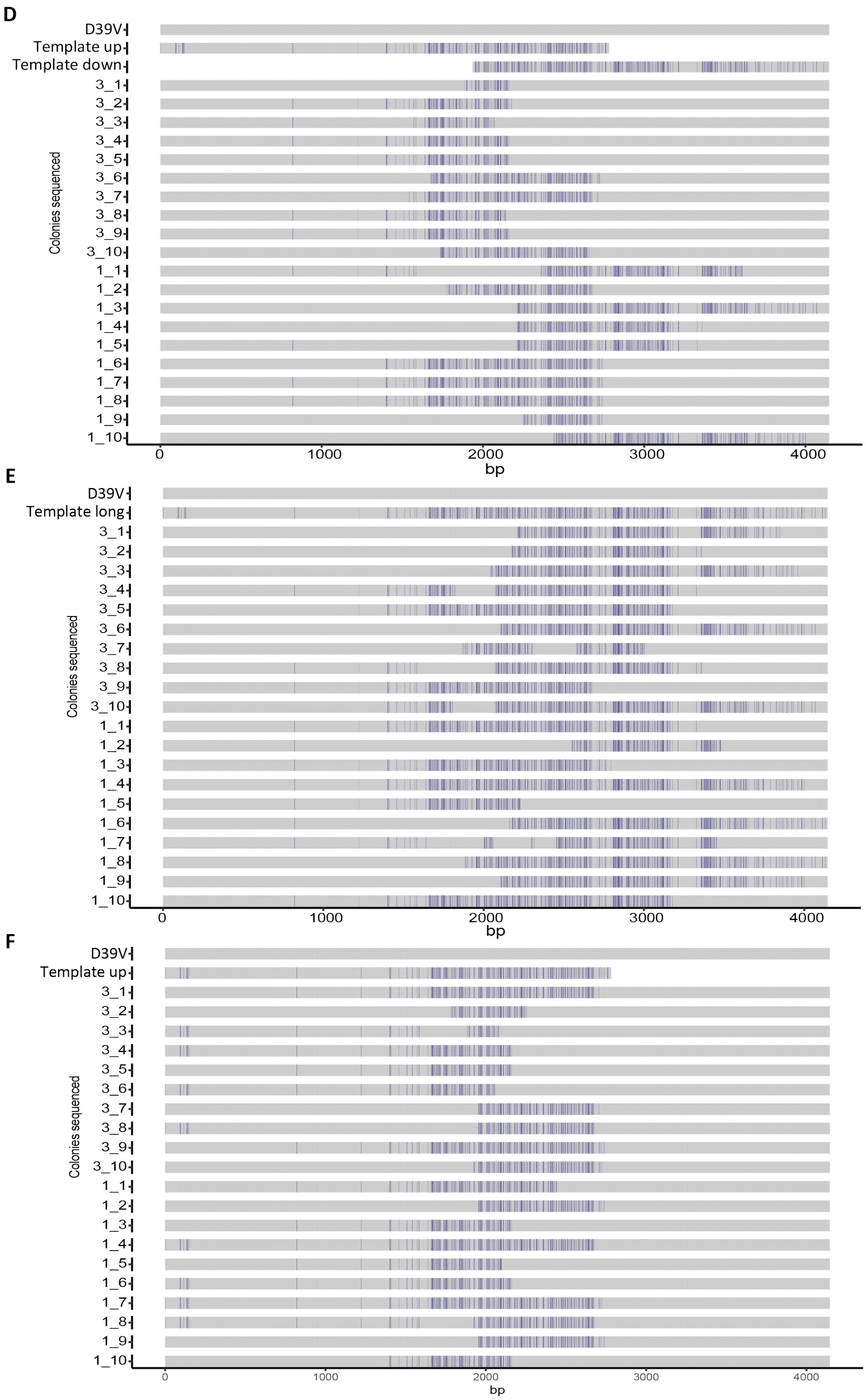
D
D39V
Template up
Template down
3_1
3_2
3_3
3_4
3_5
3_6
3_7
3_8
3_9
3_10
1_1
1_2
1_3
1_4
1_5
1_6
1_7
1_8
1_9
1_10
Colonies sequenced
0
1000
2000
3000
4000
bp
E
D39V
Template long
Colonies sequenced
bp
F
D39V
Template up
Colonies sequenced
bp

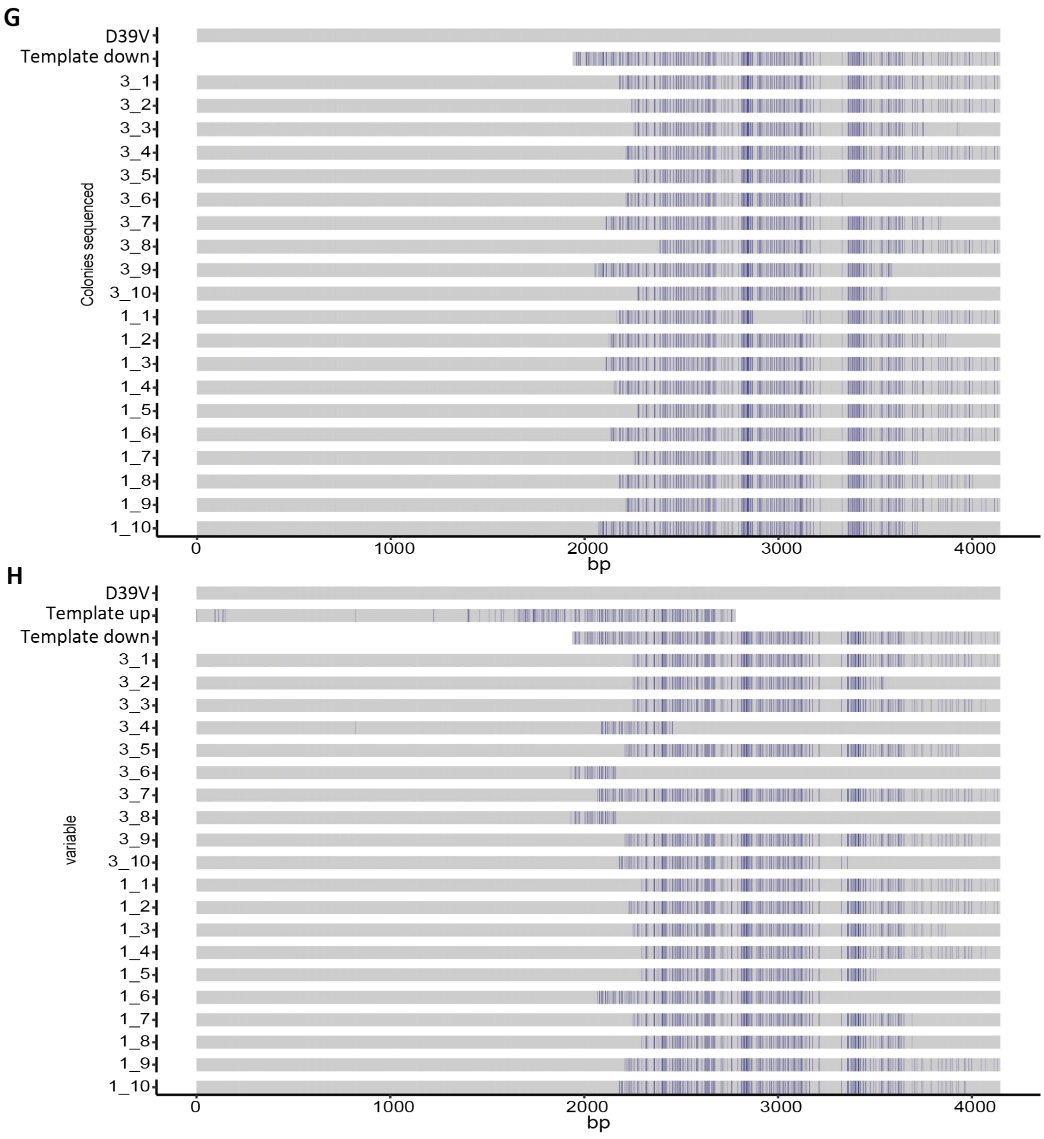


**Figure S3: Alignments of pbp2x-mraY from colonies isolated after transformation with different length and combinations of donor fragments to explain the occurrence of non-contiguous recombination.** Facets show colonies selected from different donor templates which correspond to Figure 6. Bases which match the recipient (D39V) are coloured grey, those which match the donor (11A) are shown in blue. (A) Long, (B) Up, (C) Down, (D) Up & Down, (E) Long-NH, (F) NH-Up, (G) Down-NH, and (H) NH-Up and Down-NH.

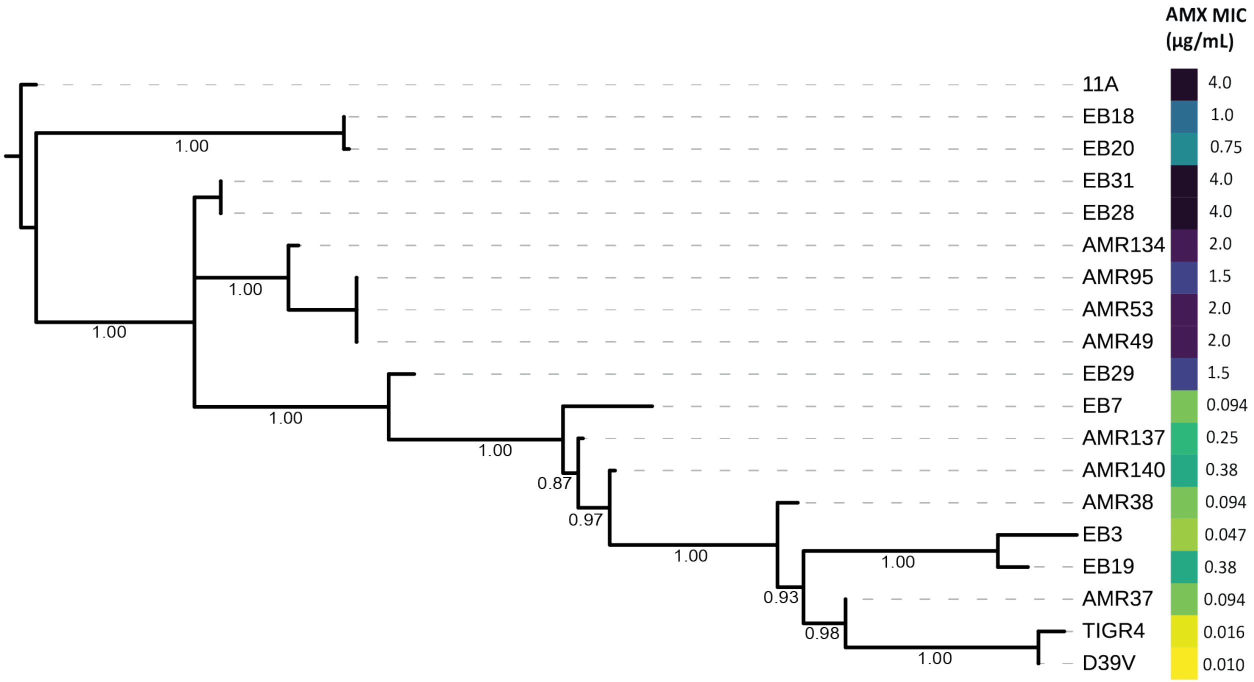


**Figure S4: Dendrogram of donor, recipient, and recombinant strains inferred from amino acid sequences of Pbp and MurM proteins**. Alignments of Pbp1a, Pbp2b, Pbp2x, and MurM amino acid sequences were used. AMX MICs are shown on the right, colour scale represents MIC and corresponds to figure 2, where yellow is an AMX MIC of 0.01 µg/mL and dark purple is an MIC of 4 µg/mL.
